# SAMPLER: Empirical distribution representations for rapid analysis of whole slide tissue images

**DOI:** 10.1101/2023.08.01.551468

**Authors:** Patience Mukashyaka, Todd B. Sheridan, Ali Foroughi pour, Jeffrey H. Chuang

## Abstract

Deep learning has revolutionized digital pathology, allowing for automatic analysis of hematoxylin and eosin (H&E) stained whole slide images (WSIs) for diverse tasks. In such analyses, WSIs are typically broken into smaller images called tiles, and a neural network backbone encodes each tile in a feature space. Many recent works have applied attention based deep learning models to aggregate tile-level features into a slide-level representation, which is then used for slide-level prediction tasks. However, training attention models is computationally intensive, necessitating hyperparameter optimization and specialized training procedures. Here, we propose SAMPLER, a fully statistical approach to generate efficient and informative WSI representations by encoding the empirical cumulative distribution functions (CDFs) of multiscale tile features. We demonstrate that SAMPLER-based classifiers are as accurate or better than state-of-the-art fully deep learning attention models for classification tasks including distinction of: subtypes of breast carcinoma (BRCA: AUC=0.911 ± 0.029); subtypes of non-small cell lung carcinoma (NSCLC: AUC=0.940±0.018); and subtypes of renal cell carcinoma (RCC: AUC=0.987±0.006). A major advantage of the SAMPLER representation is that predictive models are >100X faster compared to attention models. Histopathological review confirms that SAMPLER-identified high attention tiles contain tumor morphological features specific to the tumor type, while low attention tiles contain fibrous stroma, blood, or tissue folding artifacts. We further apply SAMPLER concepts to improve the design of attention-based neural networks, yielding a context aware multi-head attention model with increased accuracy for subtype classification within BRCA and RCC (BRCA: AUC=0.921±0.027, and RCC: AUC=0.988±0.010). Finally, we provide theoretical results identifying sufficient conditions for which SAMPLER is optimal. SAMPLER is a fast and effective approach for analyzing WSIs, with greatly improved scalability over attention methods to benefit digital pathology analysis.

## Introduction

Pathology relies on inspecting H&E-stained tissue on glass slides under a microscope to diagnose, classify, and assess different types of cancer^1^. Digital pathology enables pathologists to store, view, and analyze WSIs computationally, which has led to the development of deep learning models to automate and facilitate image analysis^2,3^,4,5. These models have achieved high accuracy in distinguishing cancerous versus non-cancerous tissue^6,7^, as well as moderate to high accuracy in identifying cancer subtypes^6,8^,9,10. Current deep learning models can also identify regions of interest in WSIs^7,11^, a valuable tool for the diagnosis process.

The large size of WSIs presents challenges for deep learning-based predictive models^6,10^,12. To deal with this issue, WSIs are usually broken into smaller images called tiles, and each tile is passed through a neural network backbone to encode each tile in a feature space^6,9^,10. Tile-level information is then aggregated to arrive at a slide-level prediction, generally using multiple instance learning (MIL) techniques. Early MIL techniques trained and quantified predictive models at the tile-level, then obtained slide-level predictions by placing equal importance on all tiles^6,7^,10. Subsequent approaches clustered tiles and concatenated cluster-level feature representations^13,14^,15. More recently, attention-based neural networks have become popular to construct slide-level representation by aggregating weighted tile-level features followed by classifying at the slide-level ^12,16^,17,18,19, e.g. via multi-head attention (MHA), hierarchical attention, dual attention, or convolutional block attention modules^12,16^,20,21,22,23. Other important approaches have included multiscale attention and vision transformer models, which utilize the correlations across tiles to improve slide-level representations ^20,24^,25.

Nonetheless, attention-based neural networks have weaknesses. Notably, they are computationally intensive because training is performed at the slide-level. This limits the range of network architectures that can be explored^26^. The need for hyperparameter optimization exacerbates this problem since it requires training a multitude of models that each need to be tested on validation data. Reliable model selection is a particular challenge for small datasets with insufficient data for a three-way data split (train/validation/test). Additionally, the variable number of tiles within WSIs requires special training procedures such as a batch size of one or a fixed number of random tiles to represent each slide. Such procedures can lead to training instability and long convergence times^27,28^. Furthermore, attention-based models tend to have performance inferior to classical weakly supervised models on external validation datasets^26^. The lack of robustness impedes reliable interpretation^29,30^ and hinders clinical utilization.

To address these challenges, we propose a fully statistical approach to construct slide-level representations from tile-level features, called “SAMpling of multiscale empirical distributions for LEarning Representations” (SAMPLER). SAMPLER is based on the concept that the distributions of tile-level deep learning features within a WSI should provide sufficient information to make clinical predictions for the WSI, even if the distributions are specified only approximately. More precisely, SAMPLER computes the empirical cumulative distribution function (CDF) of each tile-level deep learning feature, and then the quantile values of the CDFs make up the WSI’s encoded representation. Because the SAMPLER representation is based on quantile values, it does not require de novo training of a neural network. The SAMPLER approach also does not require substantial tuning of hyperparameters, other than a consideration of multiple tile sizes and the number of quantiles. These differences make SAMPLER orders of magnitude faster than attention-based models. We demonstrate that, despite its simplicity, the SAMPLER slide level representation can be used to classify cancer subtypes with accuracy comparable to attention-based deep learning models. We confirm that attention maps generated using SAMPLER correctly identify histopathologically meaningful regions, and we further show how SAMPLER can be used to improve the design of attention-based neural networks. We also find sufficient conditions for which SAMPLER is optimal.

## Results

### SAMPLER overview

We propose a statistical approach called SAMPLER to efficiently combine the feature vectors of tiles into a single WSI-level feature vector, summarized in Fig. 1 (see Methods for details). To achieve this, we consider a multi-scale tile that combines the contributions of sub-tiles of three sizes: 512 (small), 1536 (3×512, medium), and 2560 (5×512, large) pixels wide, respectively. Use of multiple sub-tile sizes provides flexibility in capturing morphological information at different scales. We employ InceptionV3 pre-trained on ImageNet as a backbone to generate a representation for each sub-tile in a 2048-dimensional feature space (Fig. 1a). Note, for brevity and following^7^, we use the term “mone” (<u>mo</u>rphological ge<u>ne</u>) to denote deep learning features that result from applying InceptionV3 to sub-tiles. Next, we encode each mone at the slide-level by specifying its decile distribution across tiles, i.e., for each mone at each sub-tile size, we report the value at which the cumulative distribution function (CDF) reaches the respective decile (Fig. 1b, c). Thus, SAMPLER represents each WSI by a 3×10×2048=61440 dimensional feature vector.

**Fig. 1:**
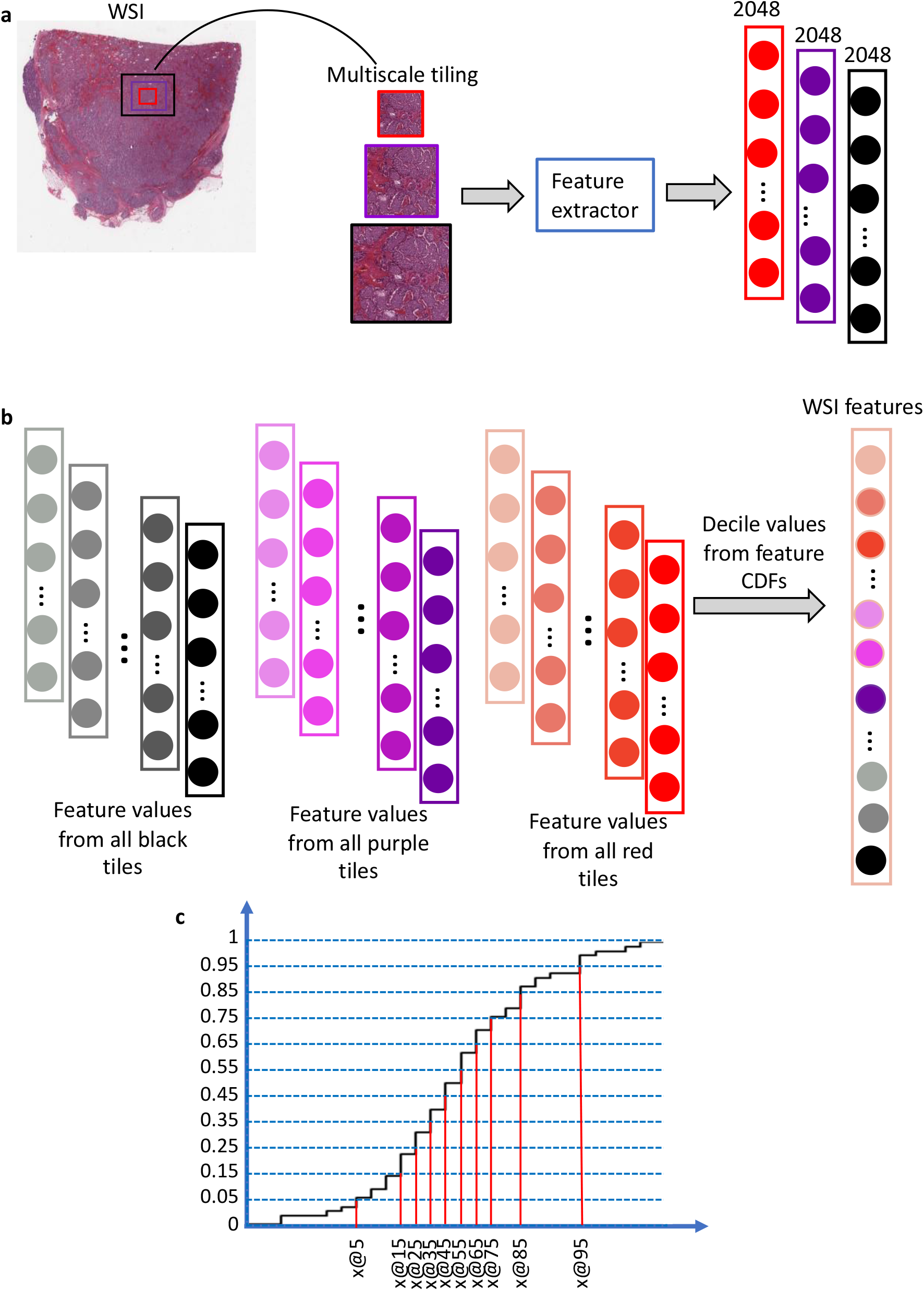
SAMPLER representation for a whole slide image. **a**. An InceptionV3 backbone is used to compute the values of deep learning features (“mones”) for each tile in a WSI. Mones are computed at 3 scales using sub-tiles shown in red, purple, and black (small, medium, large tiles respectively) outlines. **b**. For each mone, the distribution of values across tiles is computed. The decile values for each mone’s CDF, concatenated across mones and scales, form the WSI representation. **c**. Illustration of a CDF and its decile values.

### Effectiveness of SAMPLER for cancer subtype classification

To evaluate the effectiveness of SAMPLER, we examined its performance on several subtype classification tasks with a range of difficulties: distinguishing invasive ductal carcinoma (IDC) from invasive lobular carcinoma (ILC) in breast cancer (BRCA); distinguishing lung adenocarcinoma (LUAD) from lung squamous cell carcinoma (LUSC) in non-small cell lung carcinoma (NSCLC); and distinguishing clear cell (KIRC), papillary (KIRP) and chromophobe (KICH) subtypes in renal cell carcinoma (RCC). For each task, we analyzed 20X WSIs from three datasets: diagnostic formalin fixed paraffin embedded (FFPE) slides of The Cancer Genome Atlas (TCGA-DX), frozen slides of TCGA (TCGA-FR), and frozen slides of the Clinical Proteomic Tumor Analysis Consortium (CPTAC-FR) (Table 1).

**Table 1:** SAMPLER WSI level classification. AUC performance of SAMPLER cancer subtype classification for breast cancer, non-small cell lung cancer, and renal cell carcinoma. AUCs are stratified by dataset source and image magnification. Results are shown for classifiers trained using one tile scale at a time (small, medium, or large, see Figure 1) or combining all 3 tile scales (small+medium+large).

| Tissue | Dataset | Magnification | Small | Medium | Large | Small+Medium+Large |
| --- | --- | --- | --- | --- | --- | --- |
| BRCA | TCGA-DX | 20X | 0.899±0.023 | 0.898±0.035 | 0.899±0.038 | 0.911±0.029 |
| BRCA | TCGA-FR | 20X | 0.918±0.029 | 0.918±0.019 | 0.912±0.023 | 0.932±0.024 |
| BRCA | CPTAC-FR | 20X | 0.786±0.066 | 0.768±0.064 | 0.788±0.098 | 0.797±0.075 |
| NSCLC | TCGA-DX | 20X | 0.934±0.023 | 0.937±0.021 | 0.928±0.025 | 0.938±0.021 |
| NSCLC | TCGA-FR | 20X | 0.920±0.008 | 0.907±0.008 | 0.893±0.017 | 0.927±0.013 |
| NSCLC | CPTAC-FR | 20X | 0.867±0.033 | 0.868±0.031 | 0.877±0.027 | 0.883±0.027 |
| RCC | TCGA-DX | 20X | 0.984±0.004 | 0.981±0.009 | 0.982±0.007 | 0.987±0.006 |
| RCC | TCGA-FR | 20X | 0.989±0.002 | 0.984±0.003 | 0.977±0.007 | 0.991±0.002 |
| BRCA | TCGA-DX | 40X | 0.798±0.033 | 0.826±0.022 | 0.824±0.020 | 0.850±0.014 |

To train our classifiers, we implemented a two-phase pipeline. In the first phase, we used a t-test threshold on WSI-level SAMPLER features to reduce the dimensions of the data. This step improved the speed and efficiency of the subsequent classification process. In the second phase, we trained a logistic regression classifier with lasso penalty to distinguish subtypes. Our classifiers achieved statistically significant Area Under the Curve (AUC) values for all tasks with p-values of 2e-10 or lower compared to the AUC=0.5 null (Table 1). For most tasks, AUC values increased modestly when using all tile sizes (small+medium+large model) compared to when using SAMPLER features from only the small tiles (Table 1 and Extended Data Table 1). These results suggest that SAMPLER features of the 512×512 pixel tiles are sufficient for distinguishing between classes in most cases, which is consistent with previous studies^31^.

Previous studies have shown that the magnification used for WSIs can affect the accuracy of classification models^31,32^. Optimizations with respect to the magnification parameter and multiscale modeling are expected to be more impactful when models based on an individual scales perform poorly^31^. We therefore evaluated the effect of image magnification on prediction accuracy using the BRCA data, as it was a more challenging subtyping task than RCC and NSCLC subtyping. We found that multiscale modeling improved AUC (small+medium+large, AUC=0.850) compared to the single scale (small, AUC=0.798) in the 40X (p-value=0.000229). In contrast, at 20X, AUC values for all tiling approaches were high (>0.898) and the addition of multiscale had a more limited effect (small+medium+large vs small, p-value = 0.319). This indicates that the multiscale encoding approach provides more robustness to variation in magnification parameters.

### SAMPLER is as accurate as fully deep learning attention models

We next compared the performance of SAMPLER to state-of-the-art fully deep learning attention models, which we obtained from^12^ (see Methods). For the BRCA and RCC subtyping tasks, SAMPLER was the most accurate of all methods tested. For NSCLC subtyping, SAMPLER was the second-most accurate method (Table 2).

**Table 2:** AUC performance of deep learning attention models (MIL16, CLAM-SB16, DeepAttnMISL17, GCN-MIL18, DS-MIL19, HIPT12) from12 and comparison to SAMPLER.

| Architecture | BRCA Subtyping | NSCLC Subtyping | RCC Subtyping |
| --- | --- | --- | --- |
| MIL | 0.778 ± 0.091 | 0.892 ± 0.042 | 0.959 ± 0.015 |
| CLAM-SB | 0.858 ± 0.067 | 0.928 ± 0.021 | 0.973 ± 0.017 |
| <u>DeepAttnMISL</u> | 0.784 ± 0.061 | 0.778 ± 0.045 | 0.943 ± 0.016 |
| GCN-MIL | 0.840 ± 0.073 | 0.831 ± 0.034 | 0.957 ± 0.012 |
| DS-MIL | 0.838 ± 0.074 | 0.920 ± 0.024 | 0.971 ± 0.016 |
| HIPT | 0.874 ± 0.060 | **0.952 ± 0.021** | 0.980 ± 0.013 |
| **SAMPLER** | **0.911 ± 0.029** | 0.940±0.018 | **0.987±0.006** |

### SAMPLER attention maps are histopathologically meaningful

Attention maps are useful for showing which regions within a WSI are important to classification. In attention neural networks, attention layers identify tiles informative of slide labels by optimizing a weight function learned during training. The attention weights can then be used to generate attention maps showing important regions within a WSI. Although SAMPLER does not use attention layers, we can also generate analogous “attention” maps based on the differential statistics of SAMPLER mones between classes in the training data. SAMPLER attention maps differ from attention maps of fully deep learning-based models, as they are not explicitly implemented as part of the classification rule and do not have trainable parameters. Still, like attention layers of deep learning models, they output tile scores prior to classification, and tiles with large weights are most indicative of a slide-level phenotype. In principle, SAMPLER attention maps can be used as the output of a self-attention layer and replace such layers in deep learning pipelines. Briefly, we generate SAMPLER attention maps at the tile level. For each tile, we define an attention score for each mone, where the mone scores are the log-likelihood ratios of decile conditioned feature distributions. We can also generate a weighted sum of mone scores to yield a tile-level score. The weights are assigned as proportional to log-transformed p-values of a hypothesis test on the feature decile (see Methods). Scores can be further integrated across tile scales (small, medium, or large) to obtain overall attention maps (see Methods).

Through this approach, SAMPLER can yield attention maps for each tile scale and each mone, providing interpretability (Fig. 2). We observed that attention maps from different scales can be similar, but are not necessarily identical (Fig. 2a). We also noticed that the attention map that combines all scales is smoother due to its superposition of signals (Fig. 2a). This is consistent with previous studies suggesting multiscale attention maps are smoother than single scale maps^31^. Among the mones that contribute most to successful classification, some can have spatially similar attention maps, while others differ (Extended Data Figure 1, Fig. 2b). This provides evidence that different groups of mones represent distinct morphologies, each contributing separately to tissue classifications. For the slide shown in Fig 2, example tiles with high attention scores at small, medium, and large scales are shown in Fig 2c. The large tile shows irregular sheets and nests of invasive ductal carcinoma, with less intervening stroma than would be seen in invasive lobular carcinoma. Progression to medium and small tiles shows more distinctly areas of lumen formation and the cohesive nature of the cells.

**Fig. 2:**
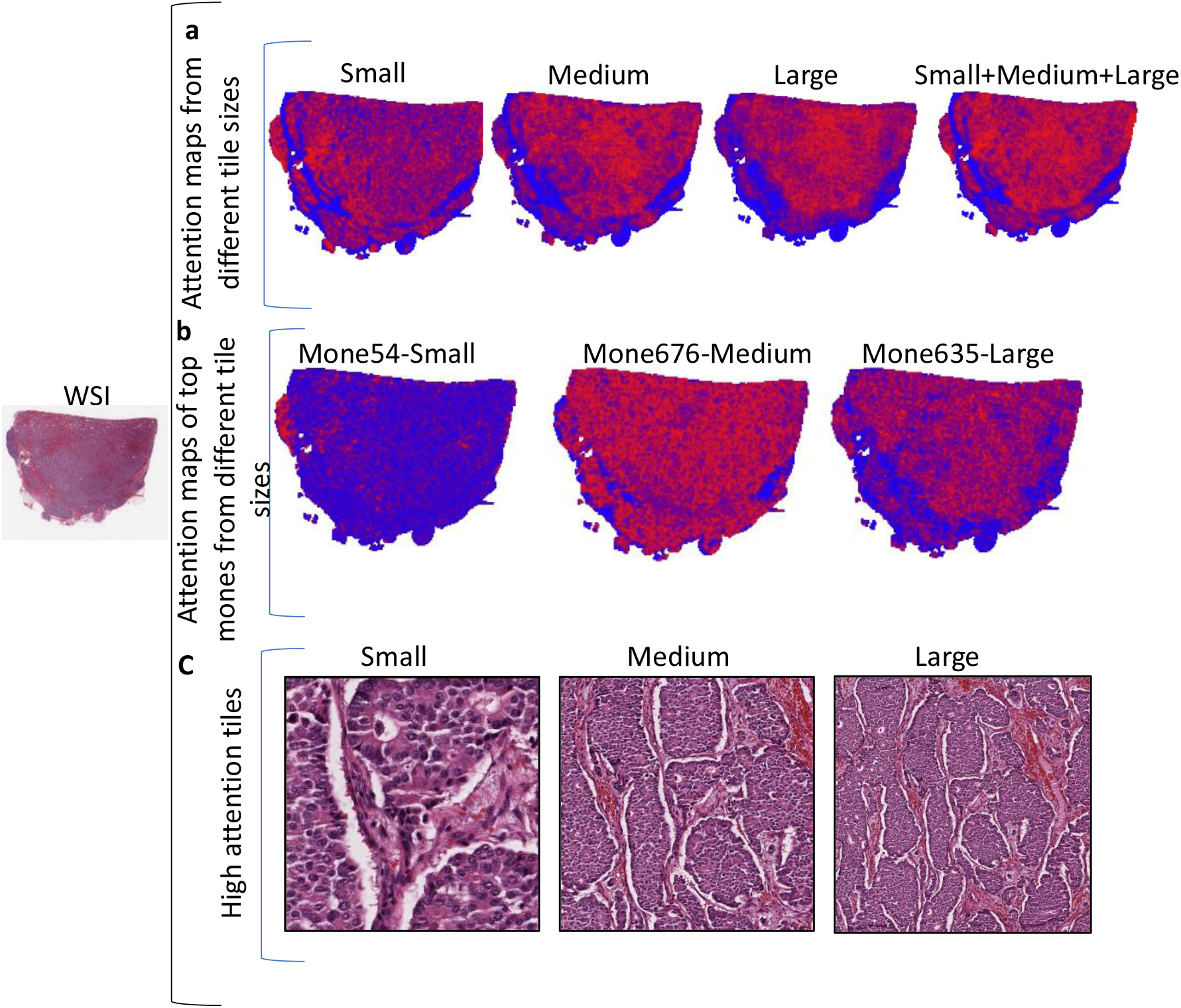
SAMPLER generates attention maps as a function of tile scale and individual features. **a**. Comparison of attention maps computed from all tile scales together (small+medium+large tiles) or individually. Red/blue coloring indicates higher/lower attention. **b**. Example attention maps for mones with high attention (top 10%). **c**. The tile with highest attention score for each of the small, medium, and large scales for the slide shown in A and B. The attention maps independently identified the same region of the slide at all three scales.

To further interpret SAMPLER-identified regions of interest, we generated attention maps for the BRCA, NSCLC, and RCC classification tasks in TCGA FFPE slides, and these were evaluated by a board-certified pathologist (Fig. 3). Tiles with high (red border) and low (blue border) attention scores were identified. Our pathologist evaluated regions with high attention scores indicative of IDC and ILC cancer subtypes (Fig. 3a), LUAD and LUSC subtypes (Fig. 3b), and KIRH, KIRC, and KIRP subtypes (Fig. 3c). Morphologies known to be associated with these subtypes were observed in tiles with high attention scores. High attention tiles for IDC showed solid sheets of tumor cells with areas of glandular lumen formation, while low attention tiles had predominantly non-neoplastic fibrous tissue with only focal areas of tumor (Fig. 3a, top panel). High attention tiles for ILC showed typical features including single cells and cords infiltrating fibrous tissue with uniform cytology, while low attention tiles showed less tumor involvement and more abundant uninvolved adipose tissue (Fig. 3a, bottom panel). LUAD high attention tiles included carcinoma with glandular lumen formation diagnostic of adenocarcinoma, while low attention tiles showed non-neoplastic lung parenchyma (Fig. 3b, top panel). LUSC high attention tiles contained solid sheets of tumor with necrosis and focal keratin pearls, while low attention tiles showed non-neoplastic lung parenchyma (Fig. 3b, bottom panel). KIRC high attention tiles showed sheets and nests of tumor cells with clear cytoplasm, distinct nuclear membranes, and arborizing vasculature, while low attention tiles contained predominantly fibrous stroma (Fig. 3c, top panel). KIRP contained tumor with papillary architecture, and low attention tiles contained non-neoplastic kidney, fibrous stroma, and areas without tissue (Fig. 3c, middle panel). KICH high attention tiles showed sheets of tumor cells with clear to granular eosinophilic cytoplasm, distinct nuclear membranes, perinuclear halos and wrinkled hyperchromatic nuclei; low attention tiles had non-neoplastic kidney and fibrous stroma (Fig. 3c, bottom panel). The high attention tiles for each tumor type contained distinctive diagnostic features for classification by the pathologist (e.g., IDC vs. ILC), supporting their utility in classification by SAMPLER. SAMPLER high attention regions were concordant with pathologist evaluations in TCGA frozen slides as well (Extended Data Figure 2). Briefly, high attention tiles contained tumor with the above features specific to the tumor type, and low attention tiles generally had areas without tissue, or contained fibrous stroma, blood, or artifacts such as tissue folds.

**Fig. 3:**
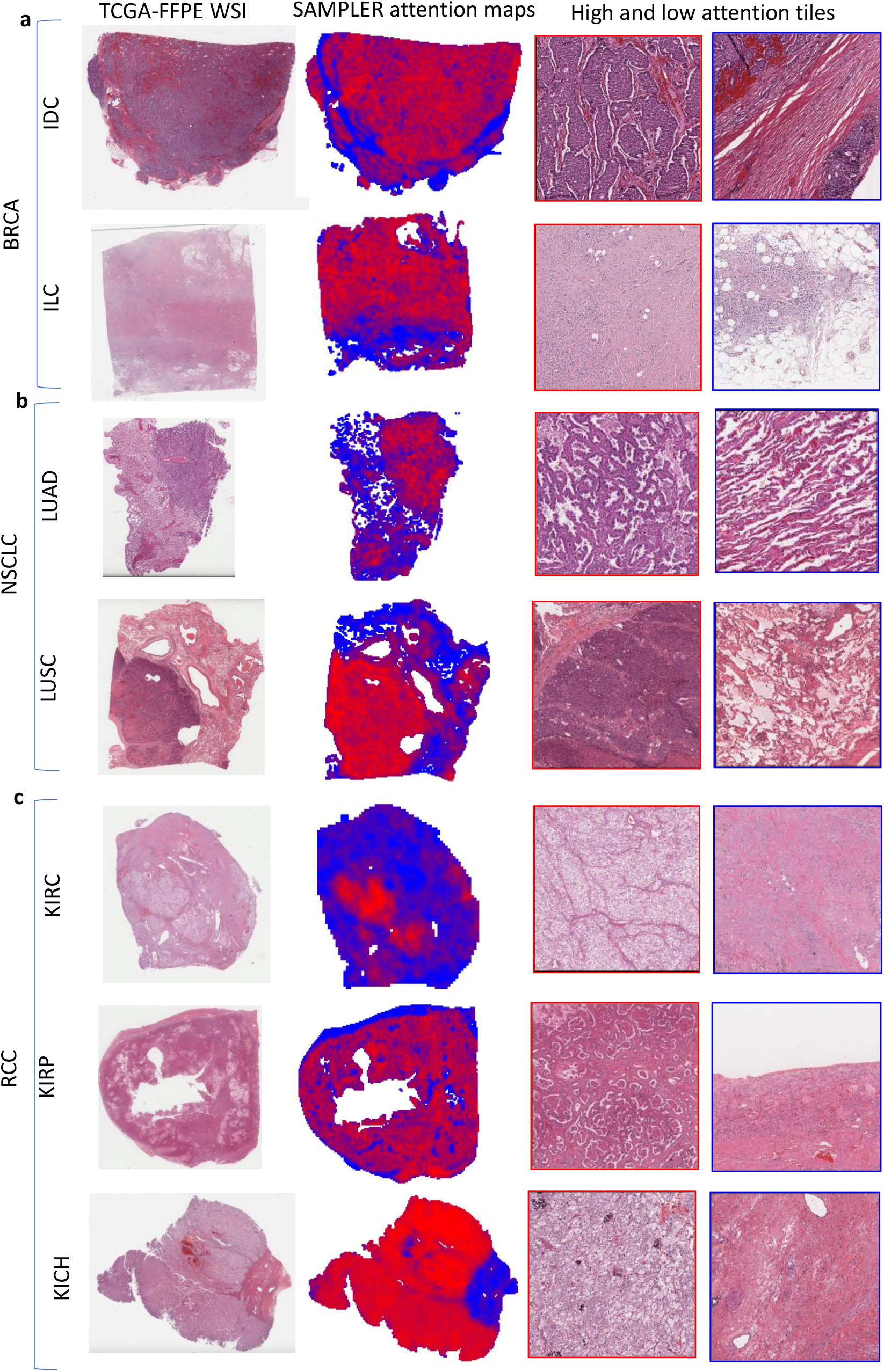
Attention maps for BRCA(a), NSCLC(b), and RCC(c) subtype classification. (Left column) WSI H&E image. (Middle column) Attention map generated by combining scores from all mones from all three tile scales. Red/blue coloring shows more/less informative regions. (Right column) Left panel indicates the highest attention tiles. Right panel indicates the lowest attention tiles. Tiles are at the 5×512 pixel scale. All examples are from TCGA-DX FFPE WSIs.

### SAMPLER classifiers are robust across data sources

Next, we tested the ability of SAMPLER-based classifiers to generalize across different data sources, i.e., TCGA and CPTAC. This was to determine robustness to batch effects in sample acquisition, staining, and scanning, which are common constraints in the applicability of H&E-based predictive models. To do this, we applied models trained on TCGA-FR to CPTAC-FR data and vice versa. This was done for both the BRCA and NSCLC subtyping tasks. The RCC task was omitted because CPTAC does not have RCC slides.

SAMPLER classifiers generalized well to external datasets, with the AUC (0.803±0.023) of the TCGA-BRCA trained model applied to CPTAC-BRCA being similar (p-value=0.812) to the internal test AUC (0.797±0.075) of classifiers trained+tested on CPTAC-BRCA (Fig. 4a). The AUC (0.792±0.027) of the CPTAC-BRCA trained model applied to TCGA-BRCA was less than (p-value=3.593e-10) the internal test AUC (0.932±0.024) of classifiers trained+tested on TCGA-BRCA (Fig. 4b). However, it was still significantly higher than the AUC=0.5 null (p-value=7.883e-18) and was similar to the internal test AUC for CPTAC-BRCA (p-value=0.845). We observed similar results for NSCLC subtyping for models trained on TCGA-FR then tested on CPTAC-FR and vice versa (Extended Data Figure 3a, b).

**Fig. 4:**
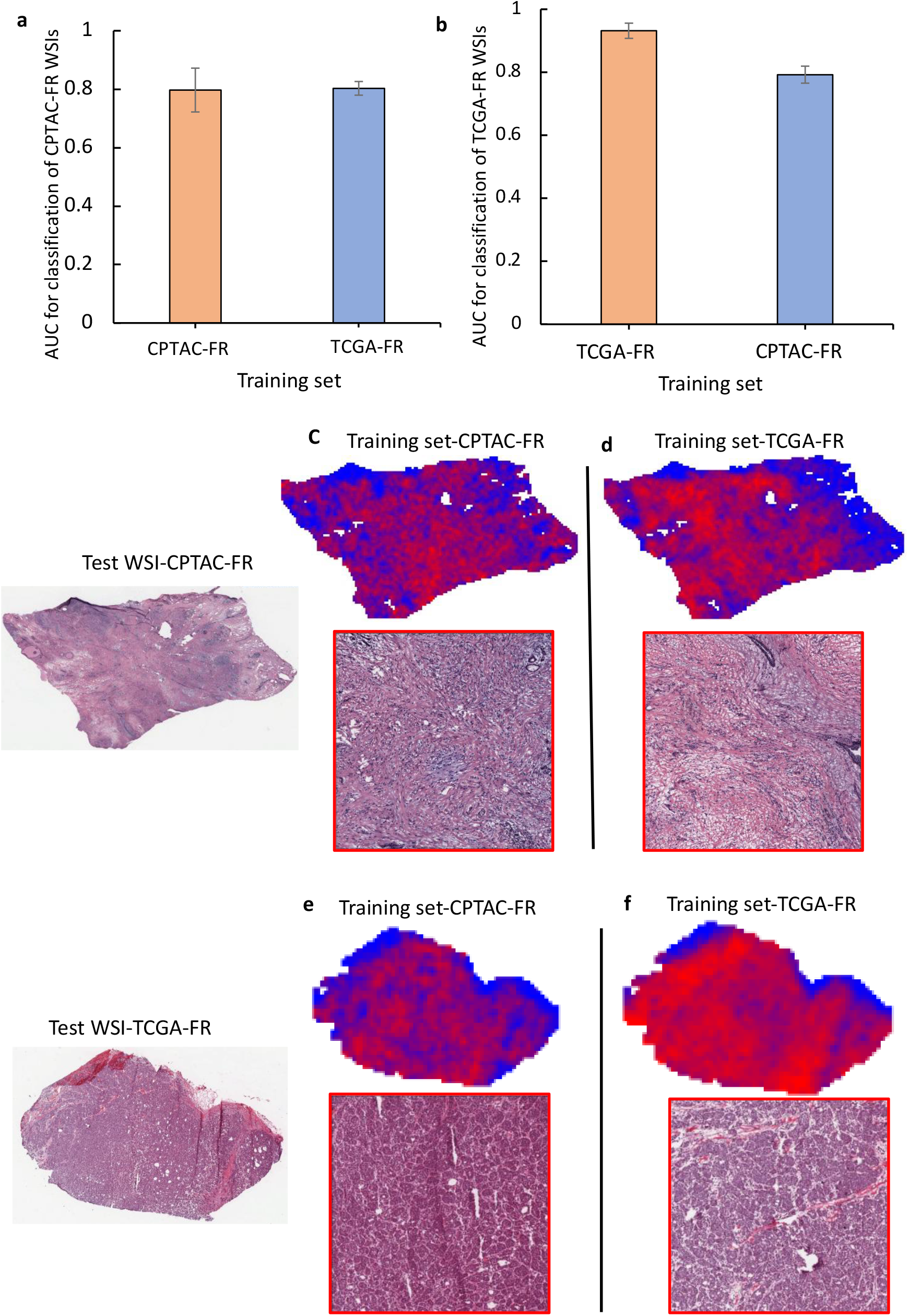
External validation on frozen WSIs. **a**. AUC and standard deviation of BRCA subtype classification of CPTAC-FR WSIs, using models trained on CPTAC-FR or TCGA-FR. **b**. AUC and standard deviation of BRCA subtype classification of TCGA-FR WSIs, using models trained on TCGA-FR or CPTAC-FR. The error bars show standard deviations from 10-fold cross-validations. **c**. Attention map and examples of high attention tiles within a CPTAC-FR WSI (ILC) for a model trained on CPTAC-FR. **d**. Attention map and examples of high attention tiles within the same CPTAC-FR WSI for a model trained on TCGA-FR. **e**. Attention map and examples of high attention tiles within a TCGA-FR WSI (IDC) for a model trained on CPTAC-FR. **f**. Attention map and examples of informative tiles within the same TCGA-FR WSI for a model trained on TCGA-FR.

Expert evaluations confirmed that regions with high attention scores in CPTAC-BRCA WSIs based on TCGA-BRCA train data were biologically meaningful (Fig. 4c, d). Similarly, based on CPTAC-BRCA train data we were able to identify biologically relevant regions in TCGA-BRCA images (Fig. 4e, f). For example, we observed infiltrating cords and single cells with low grade cytologic features in the high attention tiles in Fig. 4c and Fig. 4d, and sheets and clusters of tumor cells with areas of lumen formation in Fig. 4e and Fig. 4f. Similar observations were made in NSCLC (Extended Data Figure 3c, d). Overall, the results of the external validation and expert evaluations suggest that the classifiers and attention maps generated using SAMPLER are robust to variations across datasets.

### An optimized attention-based deep learning model based on SAMPLER

Typical attention-based deep learning models have multiple weaknesses including but not limited to the large number of trainable parameters and the fact that training at the slide-level may create training instability and overfitting in such models^26,33^. The simple effectiveness of the SAMPLER representation for WSI analysis suggests that SAMPLER concepts can guide the design of improved attention models. We expect that such improved models would have fewer parameters than typical deep attention models but maintain predictive accuracy.

We have observed several common behaviors of the SAMPLER representation. First, some SAMPLER features are systematically higher or lower between classes (Fig. 5a), i.e. individual features can have linear rather than complex differences between cancer subtypes. This suggests that a 1-layer perceptron will be sufficient for identifying important tiles, as opposed to the multi-layer perceptrons^34^ used in typical attention models. Second, different mones may correspond to different morphologies within a WSI. For example in the classification of IDC/ILC, the regions salient to mone1831 and mone1837 differ (Fig. 5b, c), and the top tiles for these mones show different morphologies (Fig. 5d,e). Specifically, mone1831 shows larger tumor clusters and darker staining compared with mone1837. This variability across mones supports the need for multi-head attention in the deep learning model. At the same time, previous studies have shown that some mones are strongly correlated and can be clustered into groups^7^. This suggests that only a small number of attention heads are needed. Lastly, multiscale classifiers are more accurate than single scale classifiers, and multiscale attention maps better distinguish tumor regions than single scale maps. This suggests that multi-scale (i.e. context aware) representations are valuable for predictive accuracy and robustness to image noise^31,32^.

**Fig. 5:**
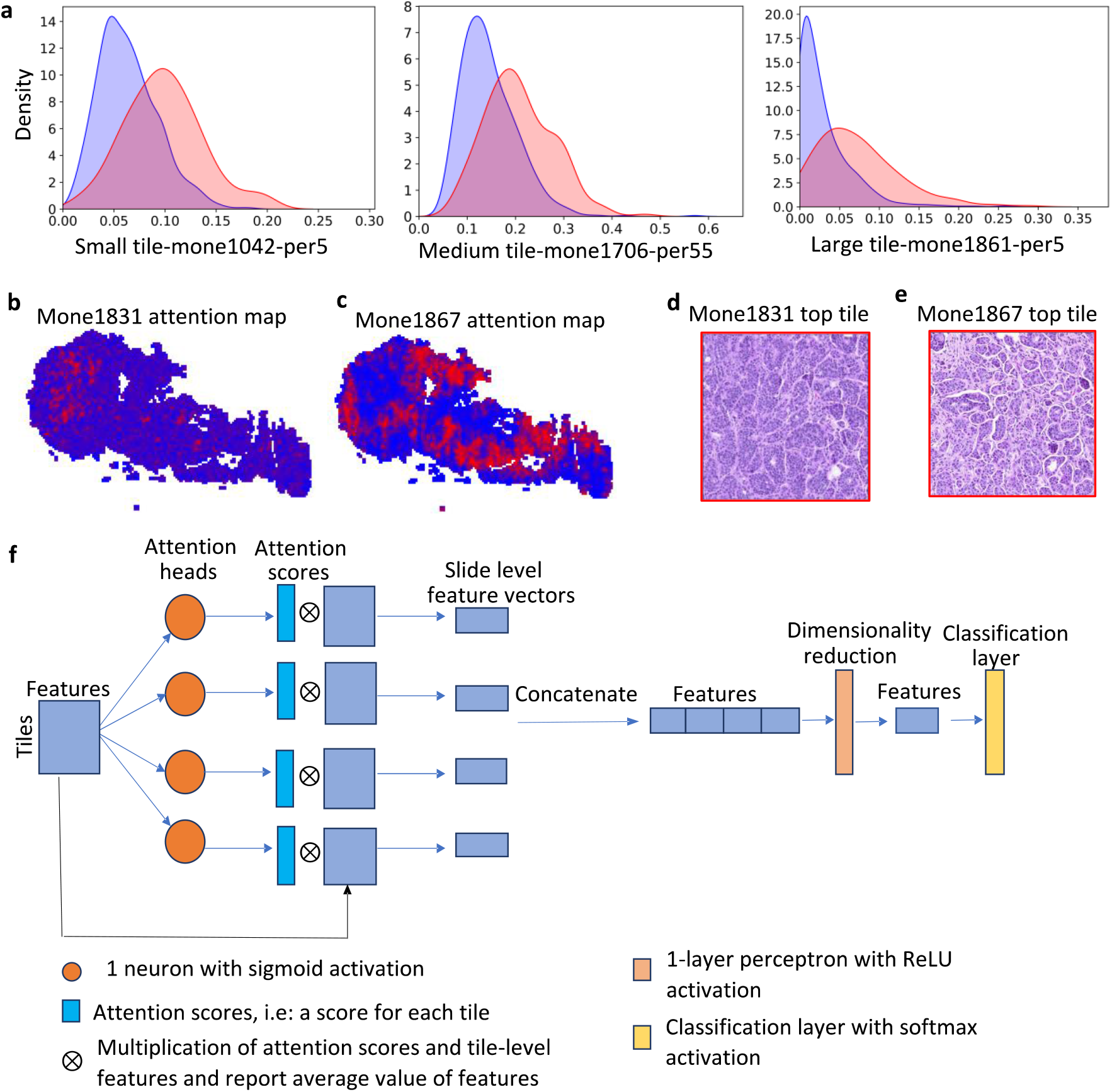
SAMPLER guides design of an improved attention model: **a**. Density plot of top SAMPLER features of each tile size for classification of IDC (red) versus ILC (blue). Features are individual mones at a specified quantile and tile size. (left) mone1042 at percentile 5 from small tiles. (middle) mone1706 at percentile 55 from medium tiles. (right) mone1861 at percentile 5 from large tiles. **b**. Attention map of mone1831 using large tiles. **c**. Attention map of mone1867 using large tiles. **d**. Example tile with high attention score from attention map in (**b**). **e**. Example tile with high attention score from attention map in (**c**). **f**. Schematic representation of context-MHA architecture.

We propose an attention deep learning model called context aware multi-head attention architecture (context-MHA) (Fig. 5f) that takes into account the behaviors observed above. The model uses a 1-layer perceptron for calculating attention, uses a small number of heads (4) for multi-head attention, and uses a context-aware representation via multi-scale tiles. These simplifications yield a much lighter network that can be more easily trained and tested than standard attention models.

We tested our context-MHA on TCGA-DX data. Compared with other deep learning models (see Table 2), context-MHA had the highest AUC for both the BRCA (AUC=0.921±0.027) and RCC (AUC=0.987±0.008) subtyping tasks. For the NSCLC subtyping task, it was second only to the HIPT model (context-MHA AUC=0.946±0.016, HIPT AUC=0.952±0.021). context-MHA also had the lowest AUC deviations across all subtyping tasks. The performance of context-MHA was also slightly better than the SAMPLER+logistic regression classifier for all tasks.

### SAMPLER is computationally efficient

A major advantage of SAMPLER over fully deep learning attention models is its low computation cost. The computation cost of SAMPLER can be divided into two parts: (1) SAMPLER WSI level feature computation, and (2) classifier training followed by testing on the validation and test sets.

To evaluate part (1), we measured the time needed to compute WSI level representations from tile level features on the TCGA-DX NSCLC dataset. When we used all the tiles in the WSI, it took less than 135 minutes (Fig. 6a) (see methods). However, SAMPLER is based on the distribution of feature values over all tiles within the WSI, and these distributions can also be approximated by sampling only a subset of tiles to further reduce the runtime (Fig. 6a) (see methods). To establish an upper bound on the runtime, for our analyses we used all tiles in each WSI unless noted. Loading tile level features from disk accounts for more than half of the computation runtime. SAMPLER is also highly parallelizable, as each WSI is processed independently. The worst-case complexity of SAMPLER is O(S × D × H × log(H)), where S, D, and H are the number of slides in the dataset, the dimensionality of the feature space, and the maximum number of sampled tiles in a WSI, respectively (Appendix). This theoretical result establishes the low computation cost of SAMPLER.

**Fig. 6:**
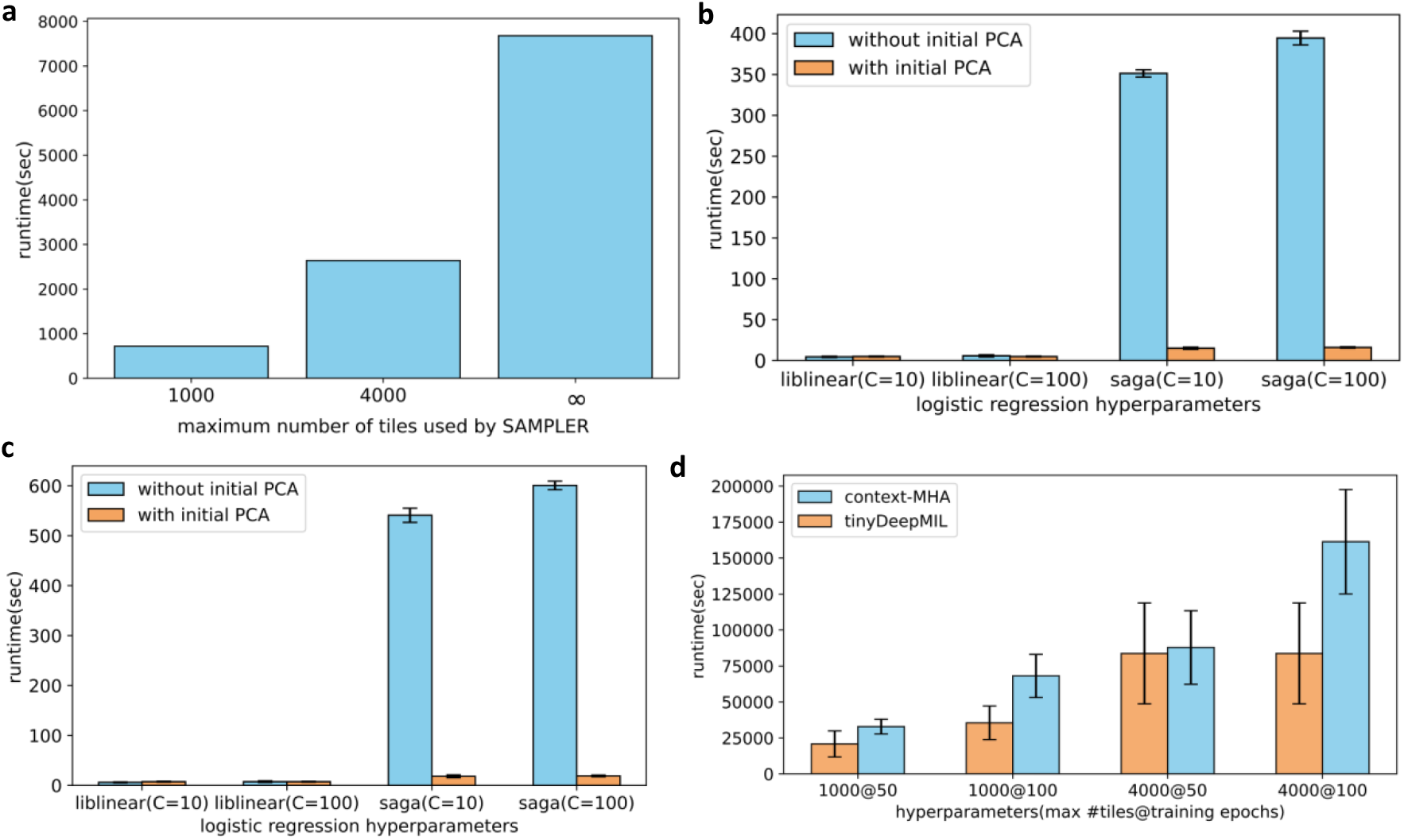
Runtime comparison of SAMPLER with attention-based deep learning models: **a**. Runtime of SAMPLER feature computation for lung WSIs of TCGA-DX, as a function of number of tiles used to calculate feature distributions. Runtimes of logistic regression models (train, validation, and test) under various hyperparameters, using **b**. small and large tiles, or **c**. small, medium, and large tiles. **d**. Runtime of context-MHA and tinyDeepMIL under various training hyperparameters using small and large tile-level features

To evaluate part (2), we measured the time needed to train, validate, and test a SAMPLER-based classifier to distinguish LUAD vs. LUSC on the TCGA-DX NSCLC dataset. We used logistic regression for classification tasks and measured runtime across different training choices. We observed that the “liblinear” optimizer was faster than “saga” using raw SAMPLER features. We also tested a model in which we simplified the SAMPLER representation by PCA analysis before inputting into the logistic regression classifier. We found that application of PCA drastically reduced runtime of “saga” (Fig. 6b, c). Interestingly, the choice between “liblinear” and “saga”, as well as the use of PCA, only had a marginal effect on the AUCs (Supplemental Files 1 and 2). For all tested parameters, PCA or “liblinear” used less than 30 seconds of runtime for training, validation, and testing (Fig. 6b, c, Supplemental Files 1 and 2).

We also compared runtimes for SAMPLER+logistic regression to runtimes of attention-based deep learning approaches. For the attention models, we analyzed our context-MHA model and a simple version of DeepMIL^34^. For DeepMIL, we specifically set its hyperparameters to minimize trainable parameters, yielding a lightweight attention model we refer to as tinyDeepMIL (see methods for details). SAMPLER+logistic regression runtimes are orders of magnitude faster than both context-MHA and tinyDeepMIL (Fig. 6d). When sampling at most 1000 tiles per WSI to construct slide-level features, SAMPLER was 454 and 288 times faster than context-MHA and tinyDeepMIL, respectively. When sampling at most 4000 tiles, SAMPLER was 332 and 316 times faster than context-MHA and tinyDeepMIL, respectively. Moreover, when using all tiles for SAMPLER and constraining context-MHA and tinyDeepMIL to use only 4000 tiles per WSI SAMPLER was still 114 and 109 times faster (Fig. 6d, methods). A key reason for this speed advantage is that SAMPLER computes WSI level representations once and saves them, while an attention-based model performs this task every time an attention layer is being trained. Therefore, hyperparameter optimization of SAMPLER-based classifiers is much faster than fully deep learning models. For example, training and testing a logistic regression classifier (solver=liblinear, max_iter=1000) is 43000 times faster than tinyDeepMIL (1000 max tiles@50epochs). Runtimes of logistic regression classifiers under additional hyperparameters are provided in Supplementary Files 1 and 2.

### SAMPLER is optimal under moderate conditions

Why does SAMPLER work, and under what conditions will it be applicable? We have found that SAMPLER is theoretically optimal for constructing slide-level representations that can recapitulate the Bayes classifier under moderate conditions. In the Appendix we prove this optimality for combining tile-level features into a slide-level representation for a given and fixed tile-level feature extractor. Although information loss in the feature extraction step cannot be recovered by SAMPLER, independence of mones and dense empirical sampling of mone CDFs are sufficient conditions to guarantee that SAMPLER can reconstruct the Bayes classifier. Moreover, the independence assumption can be relaxed for feature spaces encoding local tile interactions, and for an embedded feature space equipped with pairwise feature interactions. Two caveats are that our derivations do not provide information on the functional form of the SAMPLER-based Bayes classifier, and the optimality conditions of SAMPLER may be violated in practice. However, the strong performance of SAMPLER compared with fully deep learning models (Table 2) is likely due to two factors: (1) InceptionV3 mones are linearly associated with phenotypes^7^, and (2) empirical CDFs are smooth enough to be well-approximated by 10 deciles.

## Discussion

We have presented SAMPLER to provide a new statistical approach for aggregating tile features into WSI representations. Remarkably, SAMPLER achieves performance comparable with state-of-the-art attention-based deep learning models but is computationally cheaper by at least two orders of magnitude. SAMPLER attention maps agree with pathologist evaluations. We have shown that SAMPLER can be used to simplify and increase the accuracy of attention-based deep learning architectures. Additionally, our theoretical results provide sufficient conditions under which SAMPLER is optimal, i.e., is sufficient to recapitulate the Bayes classifier.

The core strength of SAMPLER is its statistical foundation, based on computing the quantile distributions of deep learning features across the tiles in a slide. This approach vastly simplifies the WSI into a representation with size equal to the number of deep learning features (mones) multiplied by the number of quantiles used to specify each distribution. While this statistical approach is not as flexible as attention-based neural networks, it is based on the intuitive concept that knowledge of quantile values is sufficient to distinguish tumor classes. In contrast to transformer-based approaches that suppose and attempt to account for all interactions between all tiles, SAMPLER makes the simplifying assumption that most of the relevant interactions will be local.

It is worth emphasizing SAMPLER’s advantages in computational efficiency over attention models. SAMPLER requires construction of WSI level representations only once. These representations can then be input to any downstream classifier; therefore, many classification models can be simultaneously trained and validated. Furthermore, since SAMPLER representations are at the WSI level, training using SAMPLER features is less expensive than approaches that use sets of tiles as inputs, such as attention-based deep learning models. A myriad of attention architectures have been proposed in the literature, but comparisons over architectures are often infeasible because such large models are expensive to train^26^. In contrast, we were able to design the lightweight attention model context-MHA directly from concepts motivated by SAMPLER, and it performed better than the large but naive attention models in nearly all cases.

Another strength of SAMPLER is its provision of “attention” maps with robust interpretability and that agree with pathologist evaluations. SAMPLER attention maps are robust because they are derived directly from the statistical distribution of mones (i.e. the deep learning features outputted from pre-trained architectures) and do not rely on the estimated weights of a trained classifier. Because these attention maps do not require training, they have the potential to reveal predictive regions even for WSI datasets with too few slides to train attention-based deep learning models. SAMPLER attention maps are also decomposable into the contributions of each deep learning feature, which can further assist interpretability^7^.

Our study demonstrates the efficacy of SAMPLER in generating interpretable attention maps, classifying cancer subtypes, facilitating the advancement of novel deep learning attention architectures, and reducing computational needs for slide-level classification. Still, there are important open research questions. For example, SAMPLER-related approaches may depend on the pre-trained neural network backbone used to extract mones. While we used InceptionV3 pretrained on ImageNet, it would be useful to investigate how other feature extractors work with SAMPLER. Another important question is how the number of quantiles affects accuracy and sensitivity to variations across image datasets. These and other questions may catalyze new approaches for combining tile features into slide representations to further improve tissue image analysis.

## Methods

### Dataset

We studied three cancers: Breast cancer (BRCA), Non-Small Cell Lung Carcinoma (NSCLC), and Renal Cell Carcinoma (RCC). We used two public datasets: The Cancer Genome Atlas (TCGA) and the Clinical Proteomic Tumor Analysis Consortium (CPTAC). The TCGA dataset was split into two groups: frozen (TCGA-FR) and FFPE diagnostic (TCGA-DX). All the data used from CPTAC were frozen, thus we denote them CPTAC-FR. All WSIs were processed at 20X, except for diagnostic slides of TCGA-BRCA which were processed at 20X and 40X.

### Tiling process

All WSIs described above were pre-processed, tiled, and passed through an InceptionV3 model pre-trained on ImageNet as described in^6^. We used 512 x 512 pixel tiles with 50% overlap. If a tile (small) had at least 50 % tissue, the medium (3X larger) and large (5X larger) tiles centered around the original tile were saved. Medium and large tiles were then down sampled by a factor of 3 and 5, respectively, to yield tiles 512 x 512 pixels on edge. All tiles (small, medium, and large) were then passed through InceptionV3. The 2048 features of the global average pooling (GAP) layer of InceptionV3 were written to disk.

### Train/Validation/Test splits

We used patient level splits of^12^ for TCGA-DX images. We used a 10-fold Monte Carlo cross validation for frozen slides of TCGA and CPTAC, breaking data into train (70%) and test (30%) sets at the patient level. No hyper-parameter optimization was performed on frozen slides.

### Computing slide-level representations via SAMPLER

SAMPLER generates slide-level representations by aggregating tile-level features. For each WSI, SAMPLER sorts and reports the values of each tile-level feature at several quantiles (see Appendix for precise definition). Here we used 10 quantiles (assumed M=10 in Appendix), from 5th percentile to 95 percentile in steps of 10%. Suppose X ∈ R^N,D^ is a WSI matrix where each row is a tile, and each column is a feature. For such a feature matrix, SAMPLER represents the WSI in a 10 × D dimensional feature vector.

### SAMPLER attention maps

Sample attention maps are obtained by stitching tile-level attention scores. In case of overlapping tiles, tile-level attention scores are averaged on the overlapping region. Tile-level attention scores for a binary class problem (Y ∈ {0,1}) are computed as follows: We assume the unnormalized attention score of tile T of WSI I in class y, denoted by *S*_*I*_(*T, y*), is a weighted sum of feature level attention scores:

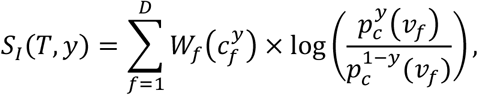

where *ν*_*f*_is the value of feature f in tile T and 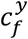 is the value of the empirical CDF of feature f for slide I at *ν*_*f*_. Let 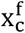 be the value of feature f when the slide-level empirical CDF of a random WSI reaches 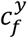. 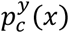 is the probability distribution function (p.d.f.) of 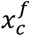 across all slides of class y. *W*_*f*_(*c*_*f*_)is the -log(p-value) of a hypothesis test. The null hypothesis assumes 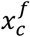 that corresponds to *c*_*f*_ does not differ between classes. We used the t-test as the hypothesis test throughout.

Given we only saved SAMPLER features at a finite number of percentiles (5th to 95th percentile in steps of 10), we used linear interpolation to compute 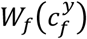 when 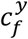 is not a multiple of 5%. Similarly, when 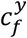 is not a multiple of 5%, we compute 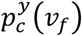 for the adjacent percentiles that are multiple of 5% and use linear interpolation. We used kernel density estimation with a Gaussian kernel to estimate the p.d.f. of 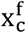. Using features belonging to one scale (small, medium, or large) generates the attention map of that scale.

For multiclass problems, to generate the attention map of a slide in class y, for each feature f we compute -log(p-value) of class y versus other classes to compute 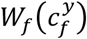. We use the average of the p.d.f. of classes ≠ *y* instead of 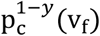.

Note: Unless otherwise stated, we use large tiles for example regions with high or low attention, as large tiles cover a larger area and provide more information regarding the region of interest.

### Logistic regression classifier training

We used a two-phase approach for classifier design. First, we used a t-test and removed features with p-values larger than a threshold (T_p_). The remaining SAMPLER features were used to train a logistic regression classifier with lasso penalty. We used a balanced class weight (class_weight= ‘balanced’). For TCGA-DX slides we tested two values of T_p_ (0.01 and 0.001), three values of C (10, 100, and 1000), two solvers (“saga” and “liblinear”), and three values for max_iter (100, 1000, 10000). We observed only minor differences in AUCs irrespective of the hyperparameters used when max_iter>100. Nonetheless, we reported the test AUC of the model with the highest validation AUC. For frozen slides we set T_p_=0.001, C=100, solver= “saga”, and max_iter=1000. We also tested if applying PCA to features that pass the t-test improves runtime and AUC. All PC dimensions were kept. For the comparison to attention deep learning models in Table 2, AUCs were obtained from^12^.

### Comparison with attention based models

We use the same data splits of^12^ borrowed from the GitHub page associated with the manuscript. The AUCs of the attention models in Table 2 are borrowed from^12^. Implementation details are provided in^12^ and the respective references for each attention model. In particular, the HIPT model uses ViT256 16 vision transformer trained for 20 epochs using Adam optimizer and a batch size of 1. Note the same splits are used to for SAMPLER-based logistic regression classifiers.

### Context aware multi-head attention (context-MHA)

The architecture of our Context-MHA is provided in (**Fig. 5f**). We used a multi-head attention model with 4 heads, each using 1-neuron with sigmoid activation as the attention module. Total attention scores of each head were normalized to 1. Attention outputs were concatenated and passed through a 1-layer perceptron using ReLU activation, reducing feature dimensionality back to the original input size. This layer was followed by the classification layer, which used softmax activation. If a slide had more than Tmax (=1000 or 4000) tiles, Tmax tiles were randomly selected. We used Adam optimizer and categorical cross entropy for training. The model with the lowest validation loss across all training epochs (=50 or 100) was saved.

### Conext-tinyDeepMIL

DeepMIL^34^ is a popular and foundational deep learning model for MIL, using a single attention mask for generating slide-level features. DeepMIL^34^ uses multi-layer perceptrons (MLPs) for assigning attention weights to tiles. In general, the attention weight of tile k is:

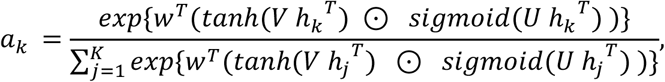

where w, U, and V denote the weights of the perceptron layers used to compute attention weights, and *h*_*k*_ is the feature vector of tile k. The number of neurons to use in each layer is a hyperparameter that can be tuned. Here, we use DeepMIL as a baseline for runtime comparisons. Therefore, we set hyperparameters of DeepMIL so that its number of trainable parameters is minimized. We used 1 neuron with tanh activation and one neuron with sigmoid activation. Therefore *w* ∈ *R* is a real number which we set to 1 for simplicity. We call this architecture context-tinyDeepMIL. context-tinyDeepMIL uses small+large tile-level representations and is hence a context-aware model. We used hyperparameters similar to context-MHA for tinyDeepMIL. Although tinyDeepMIL uses a crude attention module, its performance (BRCA: AUC=0.917 ± 0.032, NSCLC: AUC=0.936±0.017, RCC: AUC=0.984±0.011) was comparable to more complex attention models (Table 2). context-MHA and tinyDeepMIL use the same splits of^12^.

### Runtime comparison analysis

TCGA-FFPE NSCLC subtyping task was used for runtime comparisons. We considered an environment with 4 logical cores of Intel(R) Xeon(R) Gold 6150 CPUs @ 2.70GHz and 32GB of RAM on a high-performance computing (HPC) platform for all comparisons. We consider 4 parallel processes, each using 1 logical core and 8GB of RAM for computing SAMPLER features. Note SAMPLER features include all three scales (small, medium, and large). For all classifiers we compute the total runtime, i.e., the time to train a model and predict labels on validation and test WSIs across all 10 splits. We consider the small+large model for context-MHA and tinyDeepMIL. For SAMPLER-based logistic regression classifiers we consider the model with the following hyperparameters to compare runtimes with deep learning models: no PCA initialization, C=100, and solver=liblinear. For attention based deep learning models we consider runtimes of models trained for 50 epochs.

## Data and code availability

TCGA data are publicly available through the GDC potal (https://portal.gdc.cancer.gov) and CPTAC data are publicly available through the cancer imaging archive^35^ (https://www.cancerimagingarchive.net/). A minimal working code example is available at https://figshare.com/articles/software/SAMPLER_basic_code_example/23713404.

## Acknowledgement

The authors gratefully acknowledge the contribution of cyberinfrastructure high performance computing resources at the Jackson Laboratory. A.F. would like to thank the Jackson laboratory for the JAX scholar award. Research reported in this publication was supported by NIH grant R01CA230031 and NIH/NCI grant P30 CA034196.

## Author contributions

P.M conceptualized the idea, analyzed the data, and wrote the paper. T.S contributed to data analysis and provided pathological evaluations. A.F. conceptualized the idea, performed data analysis, provided the theoretical results, wrote the paper, and oversaw the work. J.C. oversaw the work, provided conceptual input, and wrote the paper.

## Extended data

**Extended Data Figure 1:**
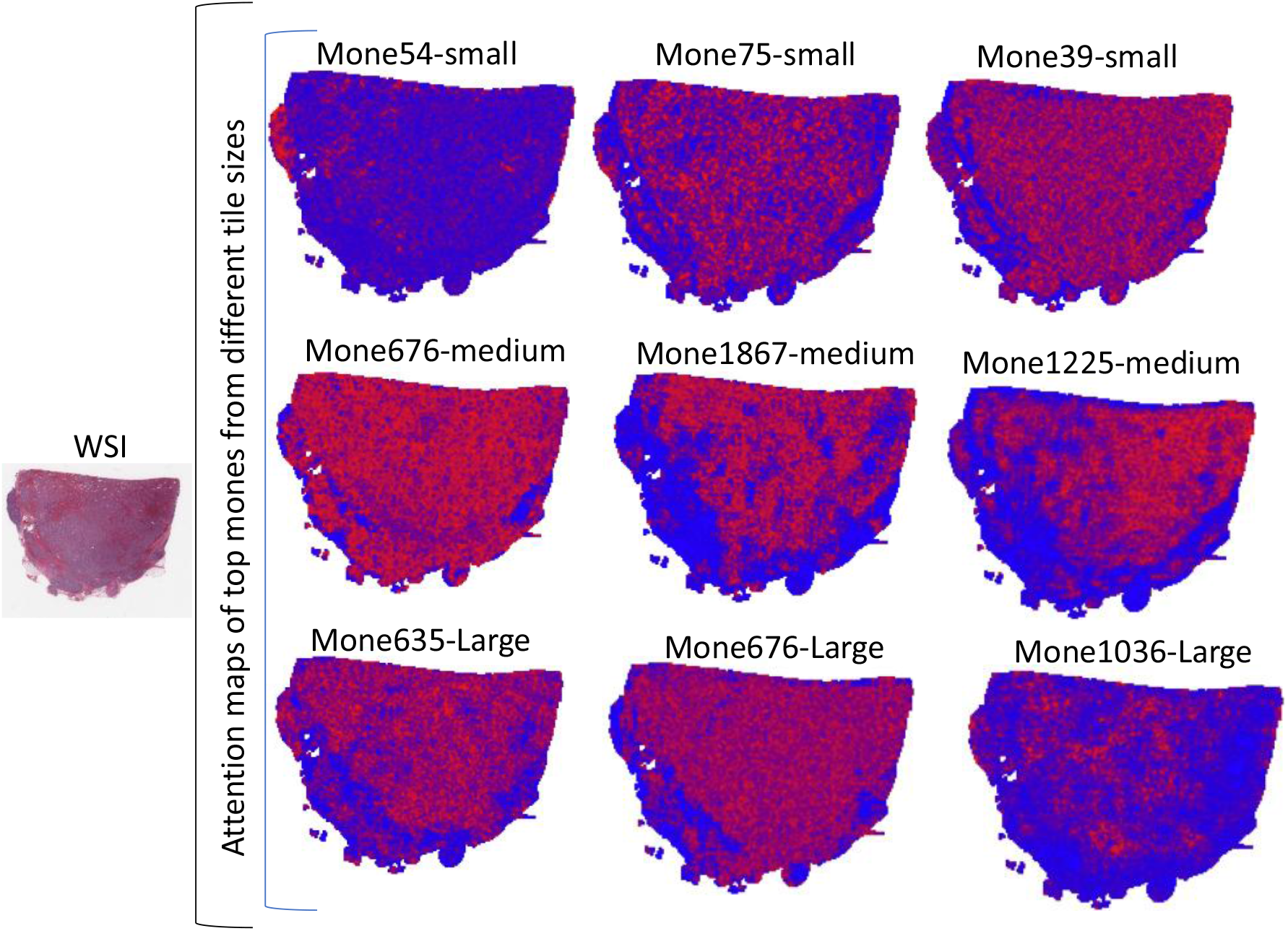
SAMPLER generates attention maps of individual mones at different scale. Example attention maps for mones in the top 10% of different tile scales: small (top panel), medium (middle panel), and large (bottom panel).

**Extended Data Figure 2:**
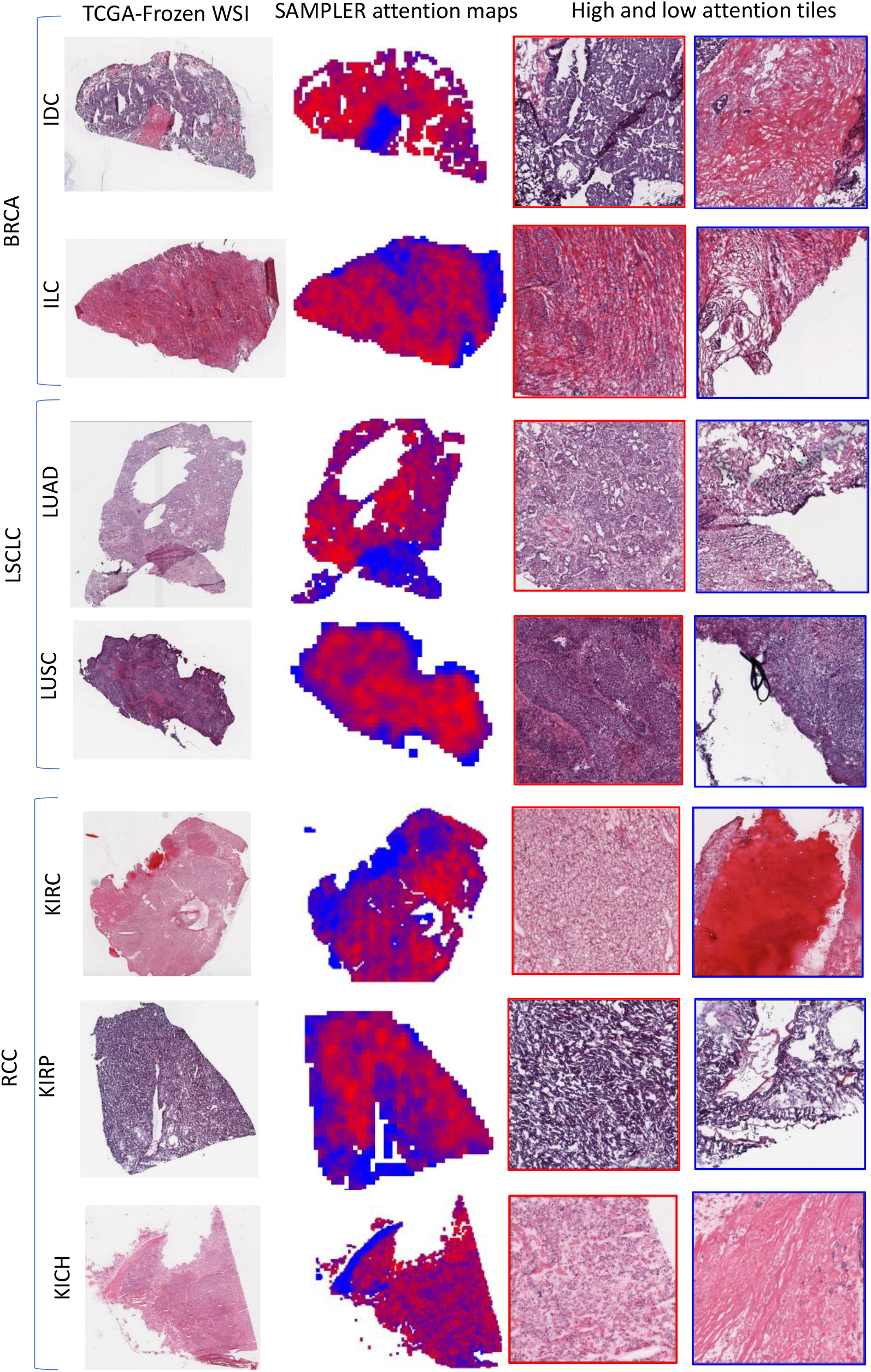
Attention maps and most and least informative tiles of BRCA, NSCLC, and RCC subtypes in TCGA-Frozen WSIs. The first column is showing the thumbnail of TCGA-FR WSIs, the second column is showing attention maps where red and blue represents informative and least informative regions respectively, and the third column is showing most informative tiles (red edges) and least informative tiles (blue edges). Top panel exhibits the BRCA subtypes IDC and ILC, middle panel exhibits the NSCLC subtypes LUAD and LUSC, and lastly the bottom panel exhibits RCC subtypes RCC, KIRC, and KIRP.

**Extended Data Figure 3:**
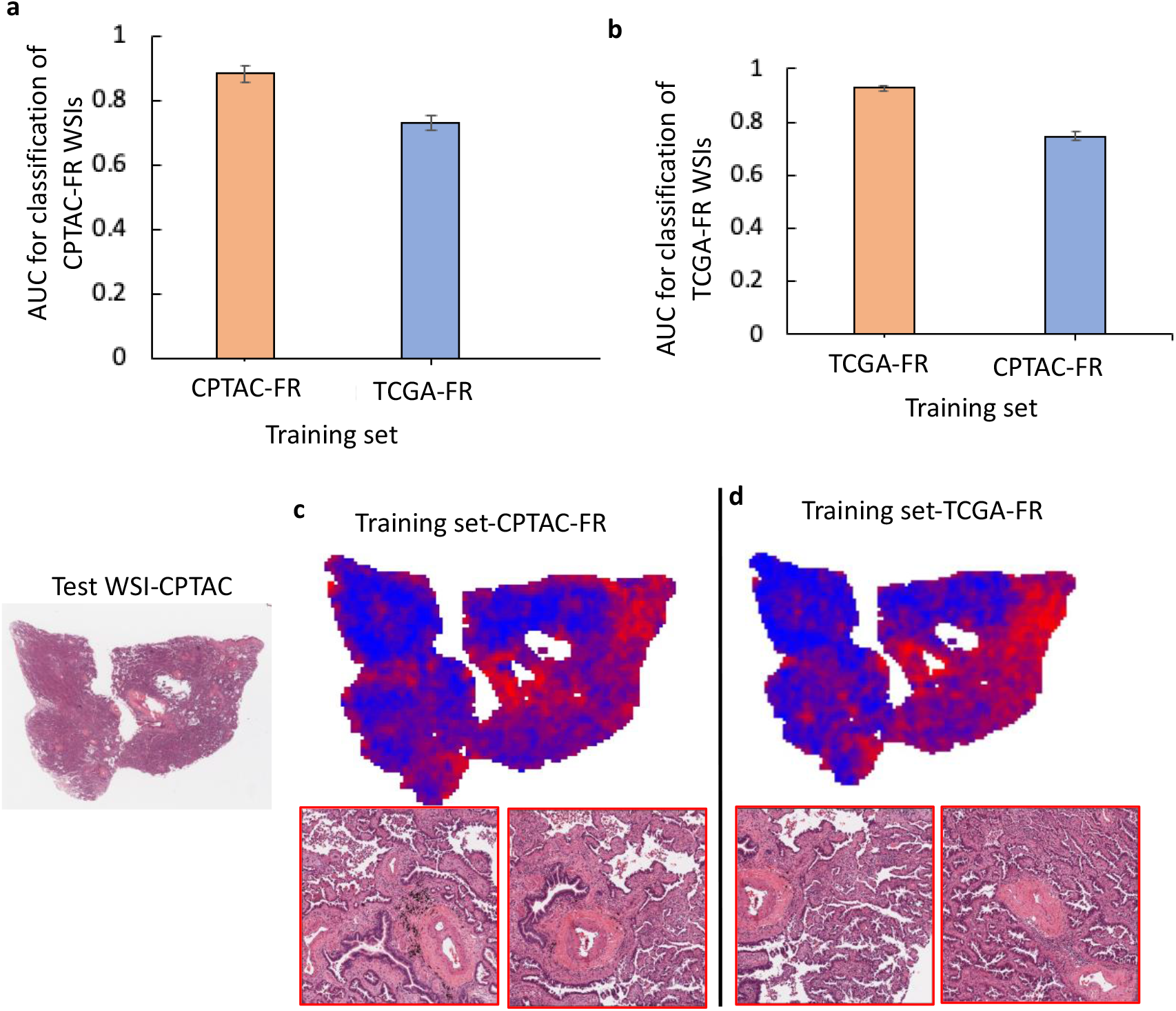
External validation of CPTAC and TCGA-FR on LSCLC. **a**. AUC and standard deviation of NSCLC subtype classification of CPTAC-FR WSIs using models trained in CPTAC and TCGA-FR (0.734±0.022). **b**. AUC and standard deviation of NSCLC subtype classification of TCGA-FR WSIs using models trained on TCGA-FR and CPTAC-FR (0.746±0.018). **c**. Attention maps and example of informative tiles of a LUAD CPTAC WSI when CPTAC training data was used, and **d**. TCGA-FR training data was used.

**Extended Data Table 1:** AUC performance of SAMPLER across different tissues, datasets for individual tile scales and their different combinations.

| Tissue | Dataset | Small | Medium | Large | Small+Medium | Small+Large | Small+Medium+Large |
| --- | --- | --- | --- | --- | --- | --- | --- |
| Breast | TCGA-DX | 0.899±0.023 | 0.898±0.035 | 0.899±0.038 | 0.907±0.026 | 0.911±0.030 | 0.911±0.029 |
| Breast | TCGA-FR | 0.918±0.029 | 0.918±0.019 | 0.912±0.023 | 0.929±0.026 | 0.929±0.025 | 0.932±0.024 |
| Breast | CPTAC-FR | 0.786±0.066 | 0.768±0.064 | 0.788±0.098 | 0.785±0.063 | 0.797±0.095 | 0.797±0.075 |
| Lung | TCGA-DX | 0.934±0.023 | 0.937±0.021 | 0.928±0.025 | 0.940±0.018 | 0.935±0.021 | 0.938±0.021 |
| Lung | TCGA-FR | 0.920±0.008 | 0.907±0.008 | 0.893±0.017 | 0.925±0.010 | 0.923±0.013 | 0.927±0.013 |
| Lung | CPTAC-FR | 0.867±0.033 | 0.868±0.031 | 0.877±0.027 | 0.876±0.030 | 0.879±0.025 | 0.883±0.027 |
| Kidney | TCGA-DX | 0.984±0.004 | 0.981±0.009 | 0.982±0.007 | 0.985±0.007 | 0.986±0.006 | 0.987±0.006 |
| Kidney | TCGA-FR | 0.798±0.033 | 0.826±0.022 | 0.824±0.020 | 0.991±0.002 | 0.991±0.002 | 0.991±0.002 |

## APPENDIX

1. **1. Complexity of SAMPLER**. SAMPLER relies on sorting all features of all tiles for each WSI. Note each slide is independently processed. As many popular sorting algorithms have a worst case time complexity of *O*(*n* log *n*), the worst case complexity of SAMPLER is *O*(*SDH* log(*H*)), where *S* is the number of WSIs in the dataset, *H* the maximum number of tiles a WSI may have, and *D* is the number of features to be sorted.
2. **2. Optimality of SAMPLER**. SAMPLER is optimal under moderate assumptions. Here we find sufficient conditions of optimality, but our conjecture is that these assumptions can be relaxed. Here we only consider a binary classification problem as generalization to multiple classes is similar.

### Definition 1.

Let *V* be a vector, *V*_*i*_ is the *i*^th^ element of *V*. Let *A* be a matrix. *A*_*i*_ is the *i*^th^ row, 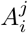 is the *j*^th^ column, and *A*^*j*^ is the element in *i*^th^ row and *j*^th^ column.

### Definition 2.

Let *V* = [*V*_1_, *V*_2_, *· · ·, V*_*n*_] be a vector. We define 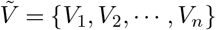. Let *A* be a matrix, *Ã*^*i*^ is the “*∼*” operation applied to *A*^*i*^.

Consider a binary classification problem with class labels *Y* = {0, 1}. Assume each tile is encoded in a *D*-dimensional feature space. Suppose *X* |*N* ∈R^*N,D*^ is a random matrix where rows represent tiles and each column is a feature. Let *C* |*N* ∈ℝ^*N*,2^ be a random matrix whose elements in row *i* are the 2D spatial coordinates of the center of the *i*^th^ tile. Let *f*_*X,C* |*Y,N*_ (*x, c*) denote the probability distribution function (p.d.f.) of random matrices in class *Y* with *N* tiles located at the coordinates in *c*. Let *f*_*N*_ (*n*) denote the probability mass function (p.m.f.) of *N*. We assume *N* and *Y* are independent.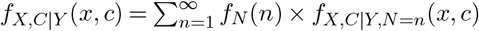. For a given fixed data point *x*, it suffices to compute the log-likelihood ratio, i.e., *L*(*x, c*) = log(*f*_*X,C Y*| =1_(*x, c*)*/f*_*X,C* |*Y* =0_(*x, c*)) to obtain the Bayes classifier. *L*(*X, C*) is the log-likelihood function applied to a random test image encoded as (*X, C*). Throughout we assume, given both *X* and *C*, the order by which tiles are placed in rows of *X* and *C* does not matter, i.e., for any row-wise permutation *P, L*(*X, C*) = *L*(*P* (*X*), *P* (*C*)). We also assume the location of the origin of the coordinate system is not important, i.e., for any translation *T, L*(*X, C*) = *L*(*X, C*^*′*^), where 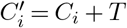.

### Definition 3.

*L*(*X, C*) is feature-decomposable if and only if 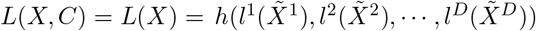 for some *h* : ℝ^*D*^ *→* ℝ and *l*^*i*^’s for almost all *X*. We have

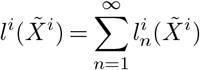

where 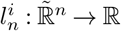 and 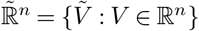. We define 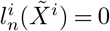 if *N* ≠ *n*.

Remark 1. Assume *L*(*X, C*) = *L*(*X*) and elements of *X* |*Y, N* are independent of each other. Then *L*(*X*) is feature-decomposable.

Remark 2. Assume 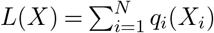 with probability 1, where each *q*_*i*_ is affine in *X*_*i*_. Then *L*(*X*) is feature-decomposable.

### Definition 4.

Let *R* ∈ℝ be a random variable. Let *Z* be the set of *K >* 0 identically distributed *R*’s. Let *S*_*Z*_(*z*) be the empirical distribution of *R* based on *Z*, i.e.,

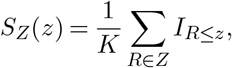

where *I*_*{}*_ is the indicator function. Fix *M*. Define *P*_*M*_ (*Z*) = [*p*_1_, …, *p*_*M*_], where, *p*_*m*_ is the smallest value of *R* ∈*Z* such that *S*_*Z*_(*R*) ≥*m/M*. *P*_*M*_ (*Z*) is the SAMPLER representation of *Z*.

### Claim 1.

Let *R* be a random variable and *Z* be the set of *K* identically distributed *R*’s with 1 ≤*K ≤H* for some *H >* 0. Assume the cumulative distribution function (CDF) of *R* is continuous and the distribution of *Z* is not degenerate. For *M* large enough, *P*_*M*_ (*Z*) is sufficient to reconstruct 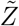 for almost all *Z*.

Proof. Observe that there exists *M* large enough, such that for all 1 ≤*K ≤H* and for each 0 ≤*k ≤K −*1, there exists 0 ≤*m ≤M −*1 such that *k/K <* (*m* + 0.5)*/M <* (*k* + 1)*/K*. Fix such *M*.

Let Ω^*H*^ be the event where all values in *Z* are distinct. Since the distribution of *R* is continuous and the distribution of *Z* is not degenerate, *P* (Ω^*H*^) = 1. Therefore, we only consider *Z* ∈Ω^*H*^. Fix *Z* ∈Ω^*H*^ with length *K*. All jumps in *Z* are of length 1*/K*. By the observation above, for each 0 ≤ *k ≤ K −*1, there exists *m* such that *p*_*m*_ is the (*k* + 1)^th^ smallest value in *Z*. Since all values in *Z* are distinct, the set of unique values in *P*_*M*_ (*Z*) is 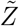. This holds for all *Z ∈* Ω^*H*^. The desired result follows.

### Theorem 1.

Assume *L*(*X, C*) is feature-decomposable. Assume for all 1 ≤*i ≤D*, there exists *s*^*i*^(.) such that 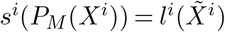 with probability 1 for some fixed *M*. Then SAMPLER features are sufficient for computing *L*(*X*) with probability 1.

Proof. Since *L*(*X, C*) is decomposable then 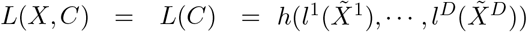 for some *h* and *l*^*i*^’s. Since *D < ∞*, then *h*(*l*^1^(*X*^1^), *· · ·, l*^*D*^(*X*^*D*^)) = *h*(*s*^1^(*P*_*M*_ (*X*^1^)), *· · ·, s*^*D*^(*P*_*M*_ (*X*^*D*^))) with probability 1.

Theorem 1 establishes the sufficient conditions for optimality of SAMPLER. The first assumption states that each feature only contributes to the Bayes classifier through its marginal distribution, i.e., higher order tile-level moments can be ignored. This may be violated in practice; however, recent work suggests linear classifiers using deep learning features enjoy accuracy comparable to complex non-linear MLPs, and are less sensitive to distribution drifts [1]. Therefore, linear models may be favored over complex MLPs given their robustness outweighs the marginal accuracy improvements of MLPs [1].

Remark 3. Let *H < ∞* be the largest number of tiles a WSI may have. By claim 1, for *M* large enough, *P*_*M*_ (*X*^*i*^) reconstructs 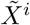. For such *M* ‘s, *s*^*i*^(*P*_*M*_ (*X*^*i*^)) = *l*^*i*^(*U* (*P*_*M*_ (*X*^*i*^))) with probability 1, where *U* (.) is the function outputting unique elements of *P*_*M*_ (*X*^*i*^).

Remark 4. If CDF’s of *X*^*i*^’s are smooth, then *P*_*M*_ (*X*^*i*^) can be a satisfactory approximation of the CDF even for small *M* ‘s. Thereby, SAMPLER-based classifiers using small *M* ‘s can still achieve high accuracy.

Remark 5. Theorem 1 establishes optimality of SAMPLER for a given and fixed feature extraction model, and optimality is only with respect to combining tile-level features into a slide-level representation. Information loss caused by the feature extractor cannot be recovered by SAMPLER.

Remark 6. Theorem 1 establishes optimality of SAMPLER by showing it is sufficient for building the Bayes classifier, but it does not comment what function family the SAMPLER-based Bayes classifier belongs to. Therefore, any off-the-shelf classifier (e.g. logistic regression, random forest, SVM) applied to SAMPLER features may be inferior to the Bayes classifier.

Feature-decomposability of the log-likelihood ratio function relies on three assumptions. (1) Only the collection of deep learning feature values affect the log-likelihood ratio, and the position of tiles within the WSI is not important. An alternate formulation is that positions of tiles do matter, but that specification of feature distributions implicitly identifies tile arrangements due to biophysical constraints. (2) Deep learning features of a tile have no interaction terms in the log-likelihood ratio function, i.e., for a tile *X*_*i*_ and two distinct features 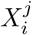 and 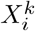, there is no term in the log-likelihood ratio function that depends on both 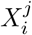 and 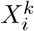. (3) Distinct deep learning features of different tiles have no interaction terms in the log-likelihood ratio function, i.e., for tiles *X*_*i*_ and *X*_*j*_, and features 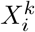 and 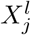 with *k* ≠ *l*, there is no term in the log-likelihood ratio function that depends on both 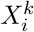 and 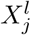. Note interaction across the same features of different tiles is allowed, i.e., for tiles *X*_*i*_ and *X*_*j*_, the interaction of features 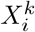 and 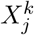 is included in 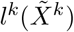. Here we relax the three assumptions above to a weaker assumption on the range of interactions across tiles.

### Definition 5.

Consider a multiple instance binary classification task, taking a set of tile features and their locations (*X* ∈ℝ^*N,D*^, *C* ∈ℝ^*N*,2^) as input. The classification task is called *locally-decomposable* if the log-likelihood ratio is feature-decomposable in the embedded feature space described below. Let 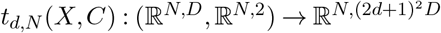 be a function that concatenates each *X*_*i*_ with the feature representation of tiles with Chebyshev distance *≤ d* to *C*_*i*_. Consider functions 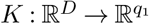 and 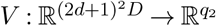 such that

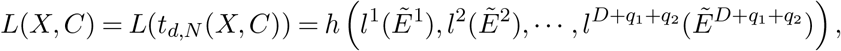

where *E* = [*X, K*(*X*), *V* (*t*_*N,d*_(*X, C*))], *K*(*X*) is a matrix whose *i*^th^ row is *K*(*X*_*i*_), *V* (*t*_*N,d*_(*X, C*)) is matrix whose *i*^th^ row is *V* (.) applied to the *i*^th^ row of 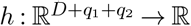, and

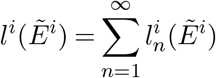

where 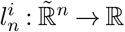. We define 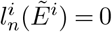 if *N* ≠ *n*.

In Definition 5, *K*(*X*) models the feature interactions within each tile. For example, *K*(*X*_*i*_) can be a polynomial kernel to account for interaction terms of the form 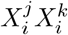. It can also model any function that operates on tile-level features, such as self-attention. *V* (*t*_*N,d*_(*X, C*)) models feature interactions across tiles that are spatially close, i.e., tiles with Chebyshev distance ≤*d* to a center tile. Therefore, it can model any function that operates on a block of adjacent tiles, e.g. attention layers that use multiple adjacent tiles to assign a weight to the center tile.

### Theorem 2.

Assume a multiple instance binary classification task on a set of images is locally-decomposable for some *d >* 0. Assume for all 1 ≤*i ≤D* + *q*_1_ + *q*_2_, there exists *s*^*i*^(.) such that 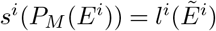 with probability 1 for some fixed *M*. Then the SAMPLER representation of *E* is sufficient for computing *L*(*X, C*) with probability 1.

Proof. Proof is analogous to theorem 1.

Remark 7. Claim 1 provides sufficient conditions for *P*_*M*_ (*Z*) to reconstruct 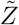 with probability 1. To apply claim 1 to *E* = [*X, K*(*X*), *V* (*t*_*N,d*_(*X, C*))] the CDFs of *K*(*X*) and *V* (*t*_*N,d*_(*X, C*)) need to be continuous in addition to *X*, upon which, for *M* large enough, SAMPLER representations would be sufficient for computing *L*(*X, C*) with probability 1 for a locally-decomposable *L*(*X, C*).

Being locally-decomposable drastically relaxes the sufficient optimality conditions of SAMPLER. For instance, polynomial *K*(*X*)’s enable modeling feature interactions within each tile. Furthermore, *K*(*X*) : ℝ^*D*^ *→* ℝ^*D*+1^ can model a self-attention layer. Consider self-attention function *a*(*x*) : ℝ^*D*^ *→* ℝ. Let

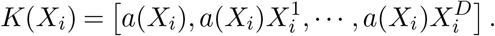

Recall *K*(*X*) is *K*(.) applied to all rows of *X*. Let *W* = [*A, W* ^1^, *· · ·, W*^*D*^] = *K*(*X*), where *A* is a column vector of all *a*(*X*_*i*_)’s and *W*^j^ is the column vector of all 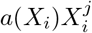’s. Let 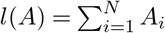 and 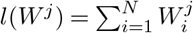. Then *l*(*W*^*j*^)*/l*(*A*) is the attention-weighted slide-level value of feature *j*. Additionally, *V* (*t*_*N,d*_(*X, C*)) models attention layers accounting for the feature interactions of spatially adjacent tiles. The *t*_*N,d*_(*X, C*) function concatenates each tile with the representations of adjacent tiles, so that their interactions can be accounted for by *V* (.), alleviating the need to keep track of tile locations. Therefore, many log-likelihood ratio functions are feature-decomposable in a higher dimensional embedding.

Remark 8. In the current manuscript, instead of training *V* (*t*_*N,d*_(*X, C*))’s from scratch, we resized medium and large tiles, and passed them through the pre-trained InceptionV3 model. Thereby, no training, i.e., parameter estimation, was performed for *V* (*t*_*N,d*_(*X, C*))’s.

### Theorem 3.

Consider a multiple instance binary classification task on a set of images with at most *H* instances per bag. Assume the CDF of *X*|*N* is continuous for all *N ≤ H*. Let *W* = *t*_*N*,1_(*X*) and 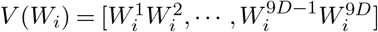. Consider the embedded feature space *E* = [*X, V* (*W*)]. Then, for *M* large enough, the SAMPLER representation of *E* is sufficient for computing *L*(*X, C*) with probability 1.

Proof. Since CDF of *X* |*N* is continuous, all elements of *X* |*N* are distinct with probability 1. Let *A*^1^ and *A*^2^ be two distinct columns of *W*. Let *A*^12^ be the column in *V* (*W*) corresponding to 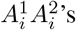. For any *a*_12_ *∈ Ã*^12^, with probability 1, there are unique *a*_1_ *∈ Ã*^1^ and *a*_2_ *∈ Ã*^2^ such that *a*_12_ = *a*_1_*a*_2_. Therefore, with probability 1, *a*_1_ and *a*_2_ belong to the same row of *W*. Iterative use of this procedure identifies the elements that comprise each row of *W* with probability 1. Thereby, we can reconstruct *W* up to a row-wise permutation. Each row of *W* identifies the relative position of the center tile with respect to its 8 adjacent tiles. Since CDF of *X* |*N* is continuous, all rows of *X* |*N* are distinct with probability 1. Therefore, with probability 1, each adjacent tile, *X*_*i,a*_, of a center tile *X*_*i*_ can be uniquely matched with another row of *W* for which *X*_*i,a*_ is the representation of the center tile. Repetitive use of this process reconstructs the location of all tile up to a translation. Since *L*(*X, C*) is invariant to row-wise permutations of *X* and *C*, and translations of *C*, the desired result follows.□

Remark 9. Assumptions of Theorem 3 can be relaxed. While Theorem 3 assumes *V* (*W*) outputs all pairwise products, fewer pairwise products are sufficient for reconstructing *W*. For example, it suffices to have 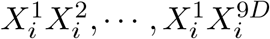. Furthermore, instead of *t*_*N*,1_ one may concatenate each *X*_*i*_ with the representation of adjacent tiles that are at the right side and at the bottom of *X*_*i*_.

Remark 10. Theorem 3 shows SAMPLER representations are optimal under moderate assumptions. Only knowledge of adjacent tile representations and pairwise feature interactions is sufficient to recapitulate all spatial information and feature interactions. While Theorem 3 provides a simple procedure to construct optimal representations, i.e., feature concatenation + pairwise interactions + large *M*, such representations may be too convoluted for a classifier to be trained on. Therefore, embeddings that are more efficient than *E* = [*X, V* (*W*)] should be explored.

