## Supplemental File 1 for "SAMPLER: Empirical distribution representations for rapid analysis of whole slide tissue images"

**Runtime and AUCs of SAMPLER without initial PCA**

| | | | | Average validation AUC | | |
|---|---|---|---|---|---|---|
| | | | | Model | | |
| Hyperparameters | Small | Medum | Large | Small-Medium | Small-Large | Small-Medium-Large |
| {'PT': 0.01, 'Miter': 100, 'solver': 'liblinear', 'C': 10} | 0.9171 | 0.929119 | 0.917847 | 0.92958 | 0.920652 | 0.927576 |
| {'PT': 0.01, 'Miter': 100, 'solver': 'liblinear', 'C': 100} | 0.915429 | 0.930925 | 0.914758 | 0.925355 | 0.922047 | 0.924074 |
| {'PT': 0.01, 'Miter': 100, 'solver': 'liblinear', 'C': 1000} | 0.915606 | 0.931429 | 0.921238 | 0.928436 | 0.927356 | 0.92535 |
| {'PT': 0.01, 'Miter': 100, 'solver': 'saga', 'C': 10} | 0.931586 | 0.933224 | 0.926848 | 0.939374 | 0.934405 | 0.936986 |
| {'PT': 0.01, 'Miter': 100, 'solver': 'saga', 'C': 100} | 0.933401 | 0.934191 | 0.928299 | 0.939512 | 0.934661 | 0.937711 |
| {'PT': 0.01, 'Miter': 100, 'solver': 'saga', 'C': 1000} | 0.933736 | 0.93438 | 0.928255 | 0.939704 | 0.934704 | 0.93786 |
| {'PT': 0.01, 'Miter': 1000, 'solver': 'liblinear', 'C': 10} | 0.917088 | 0.929851 | 0.917095 | 0.929291 | 0.91816 | 0.927999 |
| {'PT': 0.01, 'Miter': 1000, 'solver': 'liblinear', 'C': 100} | 0.915346 | 0.930683 | 0.915091 | 0.926851 | 0.920992 | 0.924741 |
| {'PT': 0.01, 'Miter': 1000, 'solver': 'liblinear', 'C': 1000} | 0.916108 | 0.931001 | 0.922147 | 0.929137 | 0.926118 | 0.926818 |
| {'PT': 0.01, 'Miter': 1000, 'solver': 'saga', 'C': 10} | 0.932865 | 0.93507 | 0.926614 | 0.936696 | 0.931636 | 0.933966 |
| {'PT': 0.01, 'Miter': 1000, 'solver': 'saga', 'C': 100} | 0.93086 | 0.935199 | 0.928109 | 0.934167 | 0.930369 | 0.932662 |
| {'PT': 0.01, 'Miter': 1000, 'solver': 'saga', 'C': 1000} | 0.930765 | 0.935296 | 0.92806 | 0.934217 | 0.930361 | 0.9328 |
| {'PT': 0.01, 'Miter': 10000, 'solver': 'liblinear', 'C': 10} | 0.916993 | 0.930188 | 0.918149 | 0.928258 | 0.91821 | 0.92709 |
| {'PT': 0.01, 'Miter': 10000, 'solver': 'liblinear', 'C': 100} | 0.915282 | 0.931401 | 0.913566 | 0.925936 | 0.919051 | 0.92507 |
| {'PT': 0.01, 'Miter': 10000, 'solver': 'liblinear', 'C': 1000} | 0.916432 | 0.930867 | 0.921054 | 0.928568 | 0.926639 | 0.927464 |
| {'PT': 0.01, 'Miter': 10000, 'solver': 'saga', 'C': 10} | 0.927106 | 0.935333 | 0.92503 | 0.934135 | 0.927108 | 0.930155 |
| {'PT': 0.01, 'Miter': 10000, 'solver': 'saga', 'C': 100} | 0.924079 | 0.934727 | 0.925524 | 0.93178 | 0.928433 | 0.930458 |
| {'PT': 0.01, 'Miter': 10000, 'solver': 'saga', 'C': 1000} | 0.923506 | 0.9352 | 0.926082 | 0.931914 | 0.928243 | 0.930695 |
| {'PT': 0.001, 'Miter': 100, 'solver': 'liblinear', 'C': 10} | 0.915015 | 0.929385 | 0.915148 | 0.930692 | 0.917332 | 0.926756 |
| {'PT': 0.001, 'Miter': 100, 'solver': 'liblinear', 'C': 100} | 0.91439 | 0.931465 | 0.915139 | 0.925984 | 0.91369 | 0.923355 |
| {'PT': 0.001, 'Miter': 100, 'solver': 'liblinear', 'C': 1000} | 0.913802 | 0.92851 | 0.921394 | 0.926835 | 0.925222 | 0.926304 |
| {'PT': 0.001, 'Miter': 100, 'solver': 'saga', 'C': 10} | 0.93051 | 0.933082 | 0.926704 | 0.938576 | 0.933879 | 0.937 |
| {'PT': 0.001, 'Miter': 100, 'solver': 'saga', 'C': 100} | 0.931899 | 0.933905 | 0.927756 | 0.939772 | 0.934184 | 0.937861 |
| {'PT': 0.001, 'Miter': 100, 'solver': 'saga', 'C': 1000} | 0.932046 | 0.933862 | 0.927858 | 0.939913 | 0.934327 | 0.93772 |
| {'PT': 0.001, 'Miter': 1000, 'solver': 'liblinear', 'C': 10} | 0.914781 | 0.930571 | 0.915568 | 0.932552 | 0.916774 | 0.929149 |
| {'PT': 0.001, 'Miter': 1000, 'solver': 'liblinear', 'C': 100} | 0.912999 | 0.931714 | 0.91704 | 0.927067 | 0.920535 | 0.924494 |
| {'PT': 0.001, 'Miter': 1000, 'solver': 'liblinear', 'C': 1000} | 0.913533 | 0.931243 | 0.921674 | 0.92706 | 0.926355 | 0.925304 |
| {'PT': 0.001, 'Miter': 1000, 'solver': 'saga', 'C': 10} | 0.930597 | 0.936568 | 0.925989 | 0.936432 | 0.93072 | 0.935054 |
| {'PT': 0.001, 'Miter': 1000, 'solver': 'saga', 'C': 100} | 0.929386 | 0.935818 | 0.928097 | 0.934244 | 0.92932 | 0.933189 |
| {'PT': 0.001, 'Miter': 1000, 'solver': 'saga', 'C': 1000} | 0.929057 | 0.935495 | 0.927863 | 0.933862 | 0.929178 | 0.932996 |
| {'PT': 0.001, 'Miter': 10000, 'solver': 'liblinear', 'C': 10} | 0.915489 | 0.93072 | 0.915285 | 0.932787 | 0.91705 | 0.928147 |
| {'PT': 0.001, 'Miter': 10000, 'solver': 'liblinear', 'C': 100} | 0.913819 | 0.931603 | 0.916919 | 0.927118 | 0.920724 | 0.924117 |
| {'PT': 0.001, 'Miter': 10000, 'solver': 'liblinear', 'C': 1000 | 0.911199 | 0.929493 | 0.92118 | 0.926776 | 0.923891 | 0.926286 |
| {'PT': 0.001, 'Miter': 10000, 'solver': 'saga', 'C': 10} | 0.924483 | 0.935866 | 0.92366 | 0.933817 | 0.925322 | 0.931885 |
| {'PT': 0.001, 'Miter': 10000, 'solver': 'saga', 'C': 100} | 0.919884 | 0.934909 | 0.92487 | 0.931403 | 0.92637 | 0.930437 |
| {'PT': 0.001, 'Miter': 10000, 'solver': 'saga', 'C': 1000} | 0.919972 | 0.935095 | 0.924636 | 0.930842 | 0.926621 | 0.930833 |

| | | | | | | |
|---|---|---|---|---|---|---|
| | | | | Average test AUC | | |
| | | | | Model | | |
| Hyperparameters | Small | Medum | Large | Small-Medium | Small-Large | Small-Medium-Large |
| {'PT': 0.01, 'Miter': 100, 'solver': 'liblinear', 'C': 10} | 0.917249 | 0.929284 | 0.917765 | 0.929823 | 0.920709 | 0.927712 |
| {'PT': 0.01, 'Miter': 100, 'solver': 'liblinear', 'C': 100} | 0.915453 | 0.931116 | 0.914671 | 0.925343 | 0.922237 | 0.924103 |
| {'PT': 0.01, 'Miter': 100, 'solver': 'liblinear', 'C': 1000} | 0.916034 | 0.931253 | 0.921202 | 0.928446 | 0.927461 | 0.925477 |
| {'PT': 0.01, 'Miter': 100, 'solver': 'saga', 'C': 10} | 0.931992 | 0.933257 | 0.926943 | 0.939392 | 0.934395 | 0.936889 |
| {'PT': 0.01, 'Miter': 100, 'solver': 'saga', 'C': 100} | 0.933784 | 0.934206 | 0.928404 | 0.939575 | 0.93463 | 0.937603 |
| {'PT': 0.01, 'Miter': 100, 'solver': 'saga', 'C': 1000} | 0.934115 | 0.934442 | 0.928357 | 0.939765 | 0.934677 | 0.937747 |
| {'PT': 0.01, 'Miter': 1000, 'solver': 'liblinear', 'C': 10} | 0.91724 | 0.930011 | 0.917011 | 0.929537 | 0.918415 | 0.928144 |
| {'PT': 0.01, 'Miter': 1000, 'solver': 'liblinear', 'C': 100} | 0.91535 | 0.930877 | 0.915007 | 0.926815 | 0.921268 | 0.924771 |
| {'PT': 0.01, 'Miter': 1000, 'solver': 'liblinear', 'C': 1000} | 0.91628 | 0.930866 | 0.922047 | 0.929153 | 0.926345 | 0.926958 |
| {'PT': 0.01, 'Miter': 1000, 'solver': 'saga', 'C': 10} | 0.933334 | 0.935089 | 0.926694 | 0.936772 | 0.931875 | 0.934072 |
| {'PT': 0.01, 'Miter': 1000, 'solver': 'saga', 'C': 100} | 0.931266 | 0.935193 | 0.928152 | 0.934276 | 0.930742 | 0.932969 |
| {'PT': 0.01, 'Miter': 1000, 'solver': 'saga', 'C': 1000} | 0.93117 | 0.93524 | 0.928102 | 0.934323 | 0.93074 | 0.933112 |
| {'PT': 0.01, 'Miter': 10000, 'solver': 'liblinear', 'C': 10} | 0.917146 | 0.930395 | 0.918054 | 0.928516 | 0.91841 | 0.927347 |
| {'PT': 0.01, 'Miter': 10000, 'solver': 'liblinear', 'C': 100} | 0.915307 | 0.931591 | 0.913477 | 0.925904 | 0.919271 | 0.9251 |
| {'PT': 0.01, 'Miter': 10000, 'solver': 'liblinear', 'C': 1000} | 0.916898 | 0.930784 | 0.921005 | 0.928589 | 0.926852 | 0.927594 |
| {'PT': 0.01, 'Miter': 10000, 'solver': 'saga', 'C': 10} | 0.927553 | 0.935433 | 0.925027 | 0.934264 | 0.927296 | 0.93033 |
| {'PT': 0.01, 'Miter': 10000, 'solver': 'saga', 'C': 100} | 0.924512 | 0.934778 | 0.925503 | 0.931823 | 0.928756 | 0.930695 |
| {'PT': 0.01, 'Miter': 10000, 'solver': 'saga', 'C': 1000} | 0.923943 | 0.935242 | 0.926062 | 0.931947 | 0.928614 | 0.930929 |
| {'PT': 0.001, 'Miter': 100, 'solver': 'liblinear', 'C': 10} | 0.915338 | 0.929411 | 0.915075 | 0.931088 | 0.917458 | 0.926992 |
| {'PT': 0.001, 'Miter': 100, 'solver': 'liblinear', 'C': 100} | 0.914737 | 0.931681 | 0.915064 | 0.925897 | 0.913815 | 0.923429 |
| {'PT': 0.001, 'Miter': 100, 'solver': 'liblinear', 'C': 1000} | 0.914324 | 0.928325 | 0.921329 | 0.926882 | 0.925538 | 0.926559 |
| {'PT': 0.001, 'Miter': 100, 'solver': 'saga', 'C': 10} | 0.930904 | 0.933123 | 0.926837 | 0.938595 | 0.933879 | 0.936884 |
| {'PT': 0.001, 'Miter': 100, 'solver': 'saga', 'C': 100} | 0.932267 | 0.933978 | 0.92789 | 0.939824 | 0.934161 | 0.937743 |
| {'PT': 0.001, 'Miter': 100, 'solver': 'saga', 'C': 1000} | 0.93241 | 0.933927 | 0.927936 | 0.939965 | 0.934304 | 0.937603 |
| {'PT': 0.001, 'Miter': 1000, 'solver': 'liblinear', 'C': 10} | 0.915104 | 0.930665 | 0.915498 | 0.932907 | 0.916885 | 0.92934 |
| {'PT': 0.001, 'Miter': 1000, 'solver': 'liblinear', 'C': 100} | 0.913351 | 0.931785 | 0.916987 | 0.927026 | 0.920731 | 0.924514 |
| {'PT': 0.001, 'Miter': 1000, 'solver': 'liblinear', 'C': 1000} | 0.91402 | 0.931019 | 0.921665 | 0.927111 | 0.926658 | 0.925385 |
| {'PT': 0.001, 'Miter': 1000, 'solver': 'saga', 'C': 10} | 0.931023 | 0.936572 | 0.926078 | 0.936643 | 0.930937 | 0.935161 |
| {'PT': 0.001, 'Miter': 1000, 'solver': 'saga', 'C': 100} | 0.92984 | 0.935812 | 0.928198 | 0.934327 | 0.929606 | 0.933407 |
| {'PT': 0.001, 'Miter': 1000, 'solver': 'saga', 'C': 1000} | 0.929503 | 0.93548 | 0.927964 | 0.933949 | 0.929461 | 0.933216 |
| {'PT': 0.001, 'Miter': 10000, 'solver': 'liblinear', 'C': 10} | 0.915806 | 0.930762 | 0.915221 | 0.933184 | 0.917314 | 0.928331 |
| {'PT': 0.001, 'Miter': 10000, 'solver': 'liblinear', 'C': 100} | 0.914172 | 0.931765 | 0.916811 | 0.927038 | 0.92091 | 0.924144 |
| {'PT': 0.001, 'Miter': 10000, 'solver': 'liblinear', 'C': 1000} | 0.911707 | 0.92946 | 0.921094 | 0.926685 | 0.924282 | 0.92647 |
| {'PT': 0.001, 'Miter': 10000, 'solver': 'saga', 'C': 10} | 0.924911 | 0.935931 | 0.923715 | 0.934 | 0.925543 | 0.932045 |
| {'PT': 0.001, 'Miter': 10000, 'solver': 'saga', 'C': 100} | 0.920253 | 0.934951 | 0.924847 | 0.931482 | 0.926686 | 0.93075 |
| {'PT': 0.001, 'Miter': 10000, 'solver': 'saga', 'C': 1000} | 0.920485 | 0.935186 | 0.924653 | 0.930912 | 0.926949 | 0.931167 |

| | | | | Average runtime (s) | | |
|---|---|---|---|---|---|---|
| | | | | Model | | |
| Hyperparameters | Small | Medum | Large | Small-Medium | Small-Large | Small-Medium-Large |
| {'PT': 0.01, 'Miter': 100, 'solver': 'liblinear', 'C': 10} | 3.71515 | 3.53172 | 3.7031 | 4.71714 | 5.04655 | 6.52901 |
| {'PT': 0.01, 'Miter': 100, 'solver': 'liblinear', 'C': 100} | 5.64889 | 4.91333 | 5.04698 | 5.926 | 5.50574 | 7.48878 |
| {'PT': 0.01, 'Miter': 100, 'solver': 'liblinear', 'C': 1000} | 5.14703 | 4.60755 | 4.91317 | 5.1374 | 5.53082 | 7.02794 |
| {'PT': 0.01, 'Miter': 100, 'solver': 'saga', 'C': 10} | 22.3211 | 26.2082 | 27.3367 | 48.3696 | 49.8038 | 75.7536 |
| {'PT': 0.01, 'Miter': 100, 'solver': 'saga', 'C': 100} | 19.4989 | 22.4812 | 23.949 | 42.0338 | 43.5125 | 66.4876 |
| {'PT': 0.01, 'Miter': 100, 'solver': 'saga', 'C': 1000} | 18.8532 | 21.8195 | 23.2045 | 40.7299 | 42.1966 | 63.8313 |
| {'PT': 0.01, 'Miter': 1000, 'solver': 'liblinear', 'C': 10} | 3.28445 | 3.35549 | 3.54568 | 4.39593 | 4.4971 | 5.87697 |
| {'PT': 0.01, 'Miter': 1000, 'solver': 'liblinear', 'C': 100} | 5.68871 | 4.63538 | 4.91966 | 5.40992 | 5.40818 | 7.01522 |
| {'PT': 0.01, 'Miter': 1000, 'solver': 'liblinear', 'C': 1000} | 5.24741 | 4.30125 | 4.23937 | 5.01534 | 4.95628 | 6.73033 |
| {'PT': 0.01, 'Miter': 1000, 'solver': 'saga', 'C': 10} | 181.302 | 213.855 | 229.514 | 396.044 | 412.112 | 624.032 |
| {'PT': 0.01, 'Miter': 1000, 'solver': 'saga', 'C': 100} | 204.748 | 234.437 | 248.11 | 441.712 | 455.37 | 697.338 |
| {'PT': 0.01, 'Miter': 1000, 'solver': 'saga', 'C': 1000} | 188.498 | 216.638 | 231.393 | 407.522 | 421.215 | 639.406 |
| {'PT': 0.01, 'Miter': 10000, 'solver': 'liblinear', 'C': 10} | 3.21203 | 3.07686 | 3.41546 | 4.47842 | 4.51438 | 5.7338 |
| {'PT': 0.01, 'Miter': 10000, 'solver': 'liblinear', 'C': 100} | 5.32812 | 4.79243 | 5.24262 | 5.66713 | 5.58075 | 6.95071 |
| {'PT': 0.01, 'Miter': 10000, 'solver': 'liblinear', 'C': 1000} | 5.15131 | 3.92758 | 4.34746 | 4.94895 | 4.76116 | 6.63706 |
| {'PT': 0.01, 'Miter': 10000, 'solver': 'saga', 'C': 10} | 596.509 | 694.449 | 989.522 | 1458.53 | 1381.43 | 2554.15 |
| {'PT': 0.01, 'Miter': 10000, 'solver': 'saga', 'C': 100} | 735.518 | 918.64 | 1148.59 | 1566.69 | 1757.06 | 2341.44 |
| {'PT': 0.01, 'Miter': 10000, 'solver': 'saga', 'C': 1000} | 626.713 | 818.525 | 947.874 | 1319.19 | 1531.54 | 1910.08 |
| {'PT': 0.001, 'Miter': 100, 'solver': 'liblinear', 'C': 10} | 3.61026 | 3.22372 | 3.64112 | 4.72107 | 4.53279 | 6.22429 |
| {'PT': 0.001, 'Miter': 100, 'solver': 'liblinear', 'C': 100} | 5.74892 | 4.76807 | 5.36826 | 5.87 | 6.08522 | 7.29596 |
| {'PT': 0.001, 'Miter': 100, 'solver': 'liblinear', 'C': 1000} | 6.59088 | 4.38766 | 4.68308 | 4.92242 | 5.15599 | 6.22825 |
| {'PT': 0.001, 'Miter': 100, 'solver': 'saga', 'C': 10} | 18.2105 | 22.3499 | 24.4272 | 41.1549 | 43.1642 | 66.6119 |
| {'PT': 0.001, 'Miter': 100, 'solver': 'saga', 'C': 100} | 15.9618 | 19.4546 | 21.3784 | 36.4692 | 37.7889 | 58.098 |
| {'PT': 0.001, 'Miter': 100, 'solver': 'saga', 'C': 1000} | 15.5575 | 18.9441 | 20.763 | 34.8214 | 36.5814 | 56.2417 |
| {'PT': 0.001, 'Miter': 1000, 'solver': 'liblinear', 'C': 10} | 3.37425 | 3.3147 | 3.55435 | 4.37468 | 4.40286 | 5.92314 |
| {'PT': 0.001, 'Miter': 1000, 'solver': 'liblinear', 'C': 100} | 6.06014 | 5.49979 | 5.4072 | 5.76102 | 5.69145 | 7.49188 |
| {'PT': 0.001, 'Miter': 1000, 'solver': 'liblinear', 'C': 1000} | 6.01913 | 4.81522 | 4.391 | 5.24625 | 5.47196 | 6.57897 |
| {'PT': 0.001, 'Miter': 1000, 'solver': 'saga', 'C': 10} | 149.492 | 187.437 | 205.848 | 335.298 | 351.416 | 540.896 |
| {'PT': 0.001, 'Miter': 1000, 'solver': 'saga', 'C': 100} | 170.596 | 205.266 | 227.993 | 383.751 | 394.666 | 600.663 |
| {'PT': 0.001, 'Miter': 1000, 'solver': 'saga', 'C': 1000} | 154.711 | 187.445 | 205.838 | 343.737 | 363.105 | 562.264 |
| {'PT': 0.001, 'Miter': 10000, 'solver': 'liblinear', 'C': 10} | 3.24809 | 3.19135 | 3.58949 | 4.30481 | 4.55424 | 6.03857 |
| {'PT': 0.001, 'Miter': 10000, 'solver': 'liblinear', 'C': 100} | 5.44411 | 4.88125 | 4.72417 | 5.44294 | 5.22416 | 6.76815 |
| {'PT': 0.001, 'Miter': 10000, 'solver': 'liblinear', 'C': 1000} | 5.62711 | 3.97886 | 4.48935 | 5.04928 | 5.09433 | 6.14808 |
| {'PT': 0.001, 'Miter': 10000, 'solver': 'saga', 'C': 10} | 622.245 | 676.023 | 764.678 | 1228.77 | 1294.05 | 2257.95 |
| {'PT': 0.001, 'Miter': 10000, 'solver': 'saga', 'C': 100} | 724.253 | 687.41 | 1046.01 | 1297.38 | 1507.31 | 2122.06 |
| {'PT': 0.001, 'Miter': 10000, 'solver': 'saga', 'C': 1000} | 592.933 | 617.234 | 891.675 | 1077.62 | 1395.44 | 1715.39 |
