## Supplemental File 2 for "SAMPLER: Empirical distribution representations for rapid analysis of whole slide tissue images"

**Runtime and AUCs of SAMPLER with initial PCA**

| | | | | Average validation AUC | | |
|---|---|---|---|---|---|---|
| | | | | Model | | |
| Hyperparameters | Small | Medum | Large | Small-Medium | Small-Large | Small-Medium-Large |
| {'PT': 0.01, 'Miter': 100, 'solver': 'liblinear', 'C': 10} | 0.91175 | 0.92879 | 0.909837 | 0.919239 | 0.917125 | 0.928745 |
| {'PT': 0.01, 'Miter': 100, 'solver': 'liblinear', 'C': 100} | 0.906263 | 0.927675 | 0.905894 | 0.917773 | 0.914454 | 0.925639 |
| {'PT': 0.01, 'Miter': 100, 'solver': 'liblinear', 'C': 1000} | 0.903626 | 0.927653 | 0.906132 | 0.920153 | 0.913938 | 0.922962 |
| {'PT': 0.01, 'Miter': 100, 'solver': 'saga', 'C': 10} | 0.934657 | 0.935578 | 0.929096 | 0.937734 | 0.933753 | 0.937286 |
| {'PT': 0.01, 'Miter': 100, 'solver': 'saga', 'C': 100} | 0.934996 | 0.935959 | 0.930628 | 0.938015 | 0.933896 | 0.936902 |
| {'PT': 0.01, 'Miter': 100, 'solver': 'saga', 'C': 1000} | 0.935471 | 0.936387 | 0.930348 | 0.938252 | 0.933847 | 0.937197 |
| {'PT': 0.01, 'Miter': 1000, 'solver': 'liblinear', 'C': 10} | 0.911797 | 0.92879 | 0.909736 | 0.919335 | 0.917125 | 0.928841 |
| {'PT': 0.01, 'Miter': 1000, 'solver': 'liblinear', 'C': 100} | 0.905926 | 0.927269 | 0.905658 | 0.917491 | 0.914696 | 0.925687 |
| {'PT': 0.01, 'Miter': 1000, 'solver': 'liblinear', 'C': 1000} | 0.903668 | 0.927572 | 0.905843 | 0.919242 | 0.913221 | 0.923101 |
| {'PT': 0.01, 'Miter': 1000, 'solver': 'saga', 'C': 10} | 0.924969 | 0.935216 | 0.923782 | 0.930824 | 0.927766 | 0.931499 |
| {'PT': 0.01, 'Miter': 1000, 'solver': 'saga', 'C': 100} | 0.924042 | 0.935601 | 0.926928 | 0.931763 | 0.927871 | 0.930291 |
| {'PT': 0.01, 'Miter': 1000, 'solver': 'saga', 'C': 1000} | 0.924182 | 0.93551 | 0.927319 | 0.932006 | 0.9281 | 0.930341 |
| {'PT': 0.01, 'Miter': 10000, 'solver': 'liblinear', 'C': 10} | 0.91175 | 0.92879 | 0.909736 | 0.91924 | 0.91722 | 0.928887 |
| {'PT': 0.01, 'Miter': 10000, 'solver': 'liblinear', 'C': 100} | 0.905978 | 0.927317 | 0.905703 | 0.917299 | 0.914601 | 0.926023 |
| {'PT': 0.01, 'Miter': 10000, 'solver': 'liblinear', 'C': 1000} | 0.904446 | 0.92732 | 0.90569 | 0.919103 | 0.913386 | 0.923046 |
| {'PT': 0.01, 'Miter': 10000, 'solver': 'saga', 'C': 10} | 0.919263 | 0.934563 | 0.920512 | 0.928057 | 0.924182 | 0.930426 |
| {'PT': 0.01, 'Miter': 10000, 'solver': 'saga', 'C': 100} | 0.918441 | 0.935394 | 0.925302 | 0.930512 | 0.926516 | 0.929462 |
| {'PT': 0.01, 'Miter': 10000, 'solver': 'saga', 'C': 1000} | 0.919099 | 0.935011 | 0.925938 | 0.931038 | 0.926563 | 0.929143 |
| {'PT': 0.001, 'Miter': 100, 'solver': 'liblinear', 'C': 10} | 0.92139 | 0.932068 | 0.919338 | 0.92363 | 0.912649 | 0.925391 |
| {'PT': 0.001, 'Miter': 100, 'solver': 'liblinear', 'C': 100} | 0.913627 | 0.929062 | 0.9161 | 0.923131 | 0.90941 | 0.923747 |
| {'PT': 0.001, 'Miter': 100, 'solver': 'liblinear', 'C': 1000} | 0.907571 | 0.928979 | 0.914189 | 0.920779 | 0.909588 | 0.916207 |
| {'PT': 0.001, 'Miter': 100, 'solver': 'saga', 'C': 10} | 0.933047 | 0.93677 | 0.928935 | 0.938046 | 0.933182 | 0.936919 |
| {'PT': 0.001, 'Miter': 100, 'solver': 'saga', 'C': 100} | 0.933465 | 0.936865 | 0.928936 | 0.937886 | 0.932981 | 0.936626 |
| {'PT': 0.001, 'Miter': 100, 'solver': 'saga', 'C': 1000} | 0.933462 | 0.937011 | 0.929511 | 0.937886 | 0.933116 | 0.93686 |
| {'PT': 0.001, 'Miter': 1000, 'solver': 'liblinear', 'C': 10} | 0.92139 | 0.932068 | 0.919385 | 0.92363 | 0.912649 | 0.925251 |
| {'PT': 0.001, 'Miter': 1000, 'solver': 'liblinear', 'C': 100} | 0.913625 | 0.928969 | 0.915816 | 0.923275 | 0.909463 | 0.923467 |
| {'PT': 0.001, 'Miter': 1000, 'solver': 'liblinear', 'C': 1000} | 0.908732 | 0.928824 | 0.913945 | 0.920745 | 0.90982 | 0.916063 |
| {'PT': 0.001, 'Miter': 1000, 'solver': 'saga', 'C': 10} | 0.923051 | 0.935201 | 0.925829 | 0.931907 | 0.924828 | 0.929059 |
| {'PT': 0.001, 'Miter': 1000, 'solver': 'saga', 'C': 100} | 0.920935 | 0.935748 | 0.925861 | 0.930699 | 0.925867 | 0.930261 |
| {'PT': 0.001, 'Miter': 1000, 'solver': 'saga', 'C': 1000} | 0.920409 | 0.935173 | 0.925428 | 0.930598 | 0.926132 | 0.930732 |
| {'PT': 0.001, 'Miter': 10000, 'solver': 'liblinear', 'C': 10} | 0.92139 | 0.932116 | 0.919385 | 0.923629 | 0.912649 | 0.925395 |
| {'PT': 0.001, 'Miter': 10000, 'solver': 'liblinear', 'C': 100} | 0.913864 | 0.92887 | 0.916006 | 0.922989 | 0.909272 | 0.923418 |
| {'PT': 0.001, 'Miter': 10000, 'solver': 'liblinear', 'C': 1000} | 0.907864 | 0.929368 | 0.913554 | 0.92063 | 0.909297 | 0.916312 |
| {'PT': 0.001, 'Miter': 10000, 'solver': 'saga', 'C': 10} | 0.920941 | 0.936015 | 0.923324 | 0.930038 | 0.9202 | 0.927404 |
| {'PT': 0.001, 'Miter': 10000, 'solver': 'saga', 'C': 100} | 0.916109 | 0.935679 | 0.924679 | 0.929476 | 0.923863 | 0.929292 |
| {'PT': 0.001, 'Miter': 10000, 'solver': 'saga', 'C': 1000} | 0.915848 | 0.935651 | 0.925011 | 0.929666 | 0.924701 | 0.929313 |

| | | | | Average test AUC | | |
|---|---|---|---|---|---|---|
| | | | | Model | | |
| Hyperparameters | Small | Medum | Large | Small-Mediu | Small-Large | Small-Medium-Large |
| {'PT': 0.01, 'Miter': 100, 'solver': 'liblinear', 'C': 10} | 0.912395 | 0.92885 | 0.91001 | 0.919315 | 0.917323 | 0.928756 |
| {'PT': 0.01, 'Miter': 100, 'solver': 'liblinear', 'C': 100} | 0.906805 | 0.927659 | 0.905942 | 0.917905 | 0.914583 | 0.925543 |
| {'PT': 0.01, 'Miter': 100, 'solver': 'liblinear', 'C': 1000} | 0.904159 | 0.927896 | 0.906391 | 0.920441 | 0.914508 | 0.923001 |
| {'PT': 0.01, 'Miter': 100, 'solver': 'saga', 'C': 10} | 0.935044 | 0.935691 | 0.929317 | 0.937791 | 0.93399 | 0.937463 |
| {'PT': 0.01, 'Miter': 100, 'solver': 'saga', 'C': 100} | 0.935374 | 0.936072 | 0.930783 | 0.938086 | 0.934134 | 0.937089 |
| {'PT': 0.01, 'Miter': 100, 'solver': 'saga', 'C': 1000} | 0.935849 | 0.936505 | 0.930502 | 0.938322 | 0.934085 | 0.937375 |
| {'PT': 0.01, 'Miter': 1000, 'solver': 'liblinear', 'C': 10} | 0.912442 | 0.92885 | 0.909917 | 0.919411 | 0.917323 | 0.928852 |
| {'PT': 0.01, 'Miter': 1000, 'solver': 'liblinear', 'C': 100} | 0.906473 | 0.927288 | 0.905798 | 0.917617 | 0.914821 | 0.925586 |
| {'PT': 0.01, 'Miter': 1000, 'solver': 'liblinear', 'C': 1000} | 0.904203 | 0.927843 | 0.906109 | 0.919489 | 0.913805 | 0.923094 |
| {'PT': 0.01, 'Miter': 1000, 'solver': 'saga', 'C': 10} | 0.925436 | 0.935299 | 0.924004 | 0.930902 | 0.928164 | 0.931747 |
| {'PT': 0.01, 'Miter': 1000, 'solver': 'saga', 'C': 100} | 0.924492 | 0.935628 | 0.926998 | 0.93184 | 0.928321 | 0.930653 |
| {'PT': 0.01, 'Miter': 1000, 'solver': 'saga', 'C': 1000} | 0.924633 | 0.935533 | 0.927333 | 0.93213 | 0.928562 | 0.930702 |
| {'PT': 0.01, 'Miter': 10000, 'solver': 'liblinear', 'C': 10} | 0.912395 | 0.92885 | 0.909917 | 0.919317 | 0.917418 | 0.928898 |
| {'PT': 0.01, 'Miter': 10000, 'solver': 'liblinear', 'C': 100} | 0.90652 | 0.927382 | 0.905847 | 0.91743 | 0.914726 | 0.92592 |
| {'PT': 0.01, 'Miter': 10000, 'solver': 'liblinear', 'C': 1000} | 0.904957 | 0.92756 | 0.905967 | 0.919393 | 0.913945 | 0.923138 |
| {'PT': 0.01, 'Miter': 10000, 'solver': 'saga', 'C': 10} | 0.91971 | 0.934649 | 0.920761 | 0.927971 | 0.924513 | 0.930588 |
| {'PT': 0.01, 'Miter': 10000, 'solver': 'saga', 'C': 100} | 0.918737 | 0.93544 | 0.925274 | 0.930551 | 0.926945 | 0.929707 |
| {'PT': 0.01, 'Miter': 10000, 'solver': 'saga', 'C': 1000} | 0.919498 | 0.935066 | 0.925872 | 0.931077 | 0.926997 | 0.929446 |
| {'PT': 0.001, 'Miter': 100, 'solver': 'liblinear', 'C': 10} | 0.921871 | 0.932132 | 0.919236 | 0.923585 | 0.912981 | 0.925478 |
| {'PT': 0.001, 'Miter': 100, 'solver': 'liblinear', 'C': 100} | 0.914008 | 0.929155 | 0.915812 | 0.923096 | 0.909627 | 0.923717 |
| {'PT': 0.001, 'Miter': 100, 'solver': 'liblinear', 'C': 1000} | 0.907952 | 0.929099 | 0.914216 | 0.920718 | 0.909974 | 0.916324 |
| {'PT': 0.001, 'Miter': 100, 'solver': 'saga', 'C': 10} | 0.933375 | 0.93689 | 0.929075 | 0.93819 | 0.933326 | 0.937038 |
| {'PT': 0.001, 'Miter': 100, 'solver': 'saga', 'C': 100} | 0.933796 | 0.936989 | 0.929077 | 0.938047 | 0.933219 | 0.936703 |
| {'PT': 0.001, 'Miter': 100, 'solver': 'saga', 'C': 1000} | 0.933795 | 0.937131 | 0.929647 | 0.938 | 0.933318 | 0.936986 |
| {'PT': 0.001, 'Miter': 1000, 'solver': 'liblinear', 'C': 10} | 0.921871 | 0.932132 | 0.919283 | 0.923586 | 0.912981 | 0.925335 |
| {'PT': 0.001, 'Miter': 1000, 'solver': 'liblinear', 'C': 100} | 0.91401 | 0.929058 | 0.91553 | 0.92324 | 0.90967 | 0.923431 |
| {'PT': 0.001, 'Miter': 1000, 'solver': 'liblinear', 'C': 1000} | 0.909102 | 0.928964 | 0.913986 | 0.920665 | 0.910077 | 0.916134 |
| {'PT': 0.001, 'Miter': 1000, 'solver': 'saga', 'C': 10} | 0.923502 | 0.935213 | 0.925833 | 0.931958 | 0.925129 | 0.929335 |
| {'PT': 0.001, 'Miter': 1000, 'solver': 'saga', 'C': 100} | 0.92141 | 0.935856 | 0.925879 | 0.930758 | 0.926228 | 0.930648 |
| {'PT': 0.001, 'Miter': 1000, 'solver': 'saga', 'C': 1000} | 0.920887 | 0.935241 | 0.925499 | 0.930671 | 0.926491 | 0.931122 |
| {'PT': 0.001, 'Miter': 10000, 'solver': 'liblinear', 'C': 10} | 0.921871 | 0.932181 | 0.919283 | 0.923584 | 0.912981 | 0.925479 |
| {'PT': 0.001, 'Miter': 10000, 'solver': 'liblinear', 'C': 100} | 0.914243 | 0.928968 | 0.915719 | 0.92295 | 0.909483 | 0.923387 |
| {'PT': 0.001, 'Miter': 10000, 'solver': 'liblinear', 'C': 1000} | 0.908237 | 0.929435 | 0.913606 | 0.920573 | 0.909598 | 0.916281 |
| {'PT': 0.001, 'Miter': 10000, 'solver': 'saga', 'C': 10} | 0.921388 | 0.936021 | 0.923268 | 0.930048 | 0.92063 | 0.927699 |
| {'PT': 0.001, 'Miter': 10000, 'solver': 'saga', 'C': 100} | 0.916479 | 0.935794 | 0.924581 | 0.929535 | 0.924249 | 0.929691 |
| {'PT': 0.001, 'Miter': 10000, 'solver': 'saga', 'C': 1000} | 0.91625 | 0.935721 | 0.924999 | 0.929728 | 0.925033 | 0.929711 |

| | | | | | | |
|---|---|---|---|---|---|---|
| | | | | Average runtime (s) | | |
| | | | | Model | | |
| Hyperparameters | Small | Medum | Large | Small-Medium | Small-Large | Small-Medium-Large |
| {'PT': 0.01, 'Miter': 100, 'solver': 'liblinear', 'C': 10} | 2.67801 | 3.24693 | 3.39855 | 5.80745 | 5.64439 | 8.61682 |
| {'PT': 0.01, 'Miter': 100, 'solver': 'liblinear', 'C': 100} | 2.64901 | 3.21795 | 3.338 | 5.73159 | 5.79276 | 8.77032 |
| {'PT': 0.01, 'Miter': 100, 'solver': 'liblinear', 'C': 1000} | 2.64634 | 3.04725 | 3.21827 | 5.60102 | 5.84013 | 8.79646 |
| {'PT': 0.01, 'Miter': 100, 'solver': 'saga', 'C': 10} | 3.56812 | 4.03506 | 4.20589 | 6.57538 | 6.79883 | 9.74876 |
| {'PT': 0.01, 'Miter': 100, 'solver': 'saga', 'C': 100} | 3.78326 | 4.26951 | 4.39798 | 6.59829 | 6.8308 | 9.78143 |
| {'PT': 0.01, 'Miter': 100, 'solver': 'saga', 'C': 1000} | 3.80176 | 4.0968 | 4.34409 | 6.40032 | 6.49287 | 9.18223 |
| {'PT': 0.01, 'Miter': 1000, 'solver': 'liblinear', 'C': 10} | 2.44235 | 2.782 | 2.92345 | 5.01478 | 5.21033 | 7.84859 |
| {'PT': 0.01, 'Miter': 1000, 'solver': 'liblinear', 'C': 100} | 2.43184 | 2.8144 | 2.87305 | 4.9518 | 5.1305 | 7.78436 |
| {'PT': 0.01, 'Miter': 1000, 'solver': 'liblinear', 'C': 1000} | 2.33985 | 2.67937 | 2.82966 | 4.88849 | 5.06848 | 7.66846 |
| {'PT': 0.01, 'Miter': 1000, 'solver': 'saga', 'C': 10} | 12.0853 | 12.4322 | 12.639 | 14.7547 | 15.0698 | 17.6579 |
| {'PT': 0.01, 'Miter': 1000, 'solver': 'saga', 'C': 100} | 13.1959 | 13.4645 | 13.7711 | 15.9278 | 16.1655 | 18.9098 |
| {'PT': 0.01, 'Miter': 1000, 'solver': 'saga', 'C': 1000} | 13.9601 | 14.4268 | 14.5788 | 16.7026 | 16.992 | 20.0812 |
| {'PT': 0.01, 'Miter': 10000, 'solver': 'liblinear', 'C': 10} | 2.36653 | 2.73529 | 2.85989 | 5.0745 | 5.09594 | 7.70221 |
| {'PT': 0.01, 'Miter': 10000, 'solver': 'liblinear', 'C': 100} | 2.39795 | 2.71541 | 2.86315 | 4.92083 | 5.18407 | 7.77896 |
| {'PT': 0.01, 'Miter': 10000, 'solver': 'liblinear', 'C': 1000} | 2.42687 | 2.73466 | 2.73699 | 4.77083 | 4.9957 | 7.51351 |
| {'PT': 0.01, 'Miter': 10000, 'solver': 'saga', 'C': 10} | 33.2868 | 30.6297 | 29.3062 | 31.7573 | 32.0783 | 32.1403 |
| {'PT': 0.01, 'Miter': 10000, 'solver': 'saga', 'C': 100} | 31.9182 | 27.9602 | 28.0225 | 29.9491 | 29.7708 | 30.3522 |
| {'PT': 0.01, 'Miter': 10000, 'solver': 'saga', 'C': 1000} | 34.8747 | 29.4329 | 31.3618 | 32.6849 | 33.0877 | 33.1193 |
| {'PT': 0.001, 'Miter': 100, 'solver': 'liblinear', 'C': 10} | 2.13101 | 2.68632 | 2.89713 | 4.66901 | 4.95772 | 7.43782 |
| {'PT': 0.001, 'Miter': 100, 'solver': 'liblinear', 'C': 100} | 2.19173 | 2.68463 | 2.96906 | 4.73431 | 4.99403 | 7.4284 |
| {'PT': 0.001, 'Miter': 100, 'solver': 'liblinear', 'C': 1000} | 2.12678 | 2.72082 | 2.96995 | 4.76044 | 5.05434 | 7.56335 |
| {'PT': 0.001, 'Miter': 100, 'solver': 'saga', 'C': 10} | 3.20895 | 3.75532 | 4.06045 | 5.97945 | 6.21357 | 8.71682 |
| {'PT': 0.001, 'Miter': 100, 'solver': 'saga', 'C': 100} | 3.25874 | 3.84766 | 4.12972 | 5.82951 | 6.16543 | 8.78621 |
| {'PT': 0.001, 'Miter': 100, 'solver': 'saga', 'C': 1000} | 3.29679 | 3.85636 | 4.09357 | 5.92753 | 6.18468 | 8.77324 |
| {'PT': 0.001, 'Miter': 1000, 'solver': 'liblinear', 'C': 10} | 2.2394 | 2.65723 | 2.89846 | 4.72122 | 4.85753 | 7.43606 |
| {'PT': 0.001, 'Miter': 1000, 'solver': 'liblinear', 'C': 100} | 2.11851 | 2.71604 | 2.855 | 4.59663 | 4.84352 | 7.33365 |
| {'PT': 0.001, 'Miter': 1000, 'solver': 'liblinear', 'C': 1000} | 2.17209 | 2.56779 | 2.78736 | 4.52928 | 4.85569 | 7.30794 |
| {'PT': 0.001, 'Miter': 1000, 'solver': 'saga', 'C': 10} | 12.0303 | 13.0876 | 12.8366 | 14.6617 | 15.0102 | 18.198 |
| {'PT': 0.001, 'Miter': 1000, 'solver': 'saga', 'C': 100} | 12.9095 | 13.5136 | 13.8223 | 15.6091 | 16.0525 | 18.8971 |
| {'PT': 0.001, 'Miter': 1000, 'solver': 'saga', 'C': 1000} | 13.738 | 14.1297 | 14.6249 | 16.1675 | 16.462 | 19.8089 |
| {'PT': 0.001, 'Miter': 10000, 'solver': 'liblinear', 'C': 10} | 2.06036 | 2.49128 | 2.78222 | 4.39895 | 4.7154 | 7.05626 |
| {'PT': 0.001, 'Miter': 10000, 'solver': 'liblinear', 'C': 100} | 2.08175 | 2.50093 | 2.80931 | 4.48516 | 4.69561 | 7.00698 |
| {'PT': 0.001, 'Miter': 10000, 'solver': 'liblinear', 'C': 1000} | 2.04549 | 2.45474 | 2.74351 | 4.40067 | 4.65252 | 6.961 |
| {'PT': 0.001, 'Miter': 10000, 'solver': 'saga', 'C': 10} | 42.2001 | 34.6513 | 31.5857 | 43.3934 | 38.7473 | 32.5645 |
| {'PT': 0.001, 'Miter': 10000, 'solver': 'saga', 'C': 100} | 37.4891 | 27.9012 | 27.5289 | 34.1802 | 33.2259 | 29.9245 |
| {'PT': 0.001, 'Miter': 10000, 'solver': 'saga', 'C': 1000} | 39.4967 | 29.635 | 28.4518 | 33.9731 | 33.5818 | 30.6966 |
